# Eukaryotic cell biology is temporally coordinated to support the energetic demands of protein homeostasis

**DOI:** 10.1101/2020.05.14.095521

**Authors:** John S. O’Neill, Nathaniel P. Hoyle, J. Brian Robertson, Rachel S. Edgar, Andrew D. Beale, Sew Y. Peak-Chew, Jason Day, Ana S. H. Costa, Christian Frezza, Helen C. Causton

**Affiliations:** MRC Laboratory of Molecular Biology, Cambridge, CB2 0QH, UK; Middle Tennessee State University, Murfreesboro, TN 37132, USA; Imperial College, London, UK; Department of Earth Sciences, University of Cambridge, Cambridge, CB2 3EQ, UK; MRC Cancer Unit, University of Cambridge, Cambridge, UK; Columbia University Medical Center, New York, NY10032, USA

## Abstract

Every aspect of yeast physiology is subject to robust temporal regulation, this becomes apparent under nutrient-limiting conditions ^1-6^ and results in biological oscillations whose function and mechanism is poorly resolved^7^. These yeast metabolic oscillations share features with circadian rhythms and typically interact with, but are independent of, the cell division cycle. Here we show that these cellular rhythms act to minimise energy expenditure by temporally restricting protein synthesis until sufficient cellular resources are present, whilst maintaining osmotic homeostasis and protein quality control. Although nutrient supply is constant, cells initially ‘sequester and store’ metabolic resources such as carbohydrates, amino acids, K^+^ and other osmolytes; which accumulate *via* increased synthesis, transport, autophagy and biomolecular condensation that is stimulated by low glucose and cytosolic acidification. Replete stores trigger increased H^+^ export to elevate cytosolic pH, thereby stimulating TORC1 and liberating proteasomes, ribosomes, chaperones and metabolic enzymes from non-membrane bound compartments. This facilitates a burst of increased protein synthesis, the liquidation of storage carbohydrates to sustain higher respiration rates and increased ATP turnover, and the export of osmolytes to maintain osmotic potential. As the duration of translational bursting is determined by cell-intrinsic factors, the period of oscillation is determined by the time cells take to store sufficient resources to license passage through the pH-dependent metabolic checkpoint that initiates translational bursting. We propose that dynamic regulation of ion transport and metabolic plasticity are required to maintain osmotic and protein homeostasis during remodelling of eukaryotic proteomes, and that bioenergetic constraints have selected for temporal organisation that promotes oscillatory behaviour.

## Introduction

*Saccharomyces cerevisiae* undergo oscillations in oxygen consumption and many other cellular processes that synchronise spontaneously when cells are grown at high density in aerobic, nutrient-limited culture at constant pH^1,4,6^. These metabolic cycles are thought to occur cell-autonomously, and to synchronise when the extracellular nutrient supply is insufficient to support exponential growth^7,8^. Under normal conditions, DNA replication does not occur when oxygen consumption is high, whereas respiratory rate does not affect the timing of mitosis (Extended Data Fig. 1). Thus yeast respiratory oscillations (YROs) are a population-level phenomenon that is distinct from, and occurs with a different frequency to, the cell division cycle but couples with it by imposing metabolic checkpoints on cell cycle progression^9^. Despite the importance of these oscillations, the mechanism and utility of the YRO is poorly understood^2^.

As with circadian and ultradian (<24h) rhythms in mammalian cells, YROs are accompanied by large-scale changes in transcription and metabolism, as well as marked changes in the rate of cell growth ^5,10-14^. YROs share many other key features with circadian rhythms in mammalian cells, but have shorter periods that are acutely sensitive to nutrient availability ^2,6,8^. For both oscillations, current understanding of the critical causal relationships and rate constants that function over the course of each cycle is inadequate to explain how the period of oscillation is determined^15^.

Oxygen consumption rates (OCR) across the YRO can be interpolated by measuring dissolved oxygen in continuous culture, where phases of higher oxygen consumption (HOC; OCR increases, DNA replication restricted) are distinguished from lower oxygen consumption (LOC) during the rest of the cycle (Fig. 1a, Extended Data Fig. 1). Nutrient availability was manipulated by changing the rate at which medium flowed through the culture (dilution rate); higher dilution rates increase nutrient availability, medium turnover and removal of cells ^6^. We noticed that while the period of oscillation and LOC duration lengthened as dilution rate decreased, the duration of HOC was invariant (Fig. 1a,b, Extended Data Fig. 1) suggesting that HOC and LOC are regulated by cell-intrinsic and extrinsic factors, respectively. To explain how YROs are generated, their physiological consequences, and to identify factors that determine oscillatory period, we sought to understand the differential activities occurring during LOC versus HOC, and at the transition between these two states.

**Figure 1.**
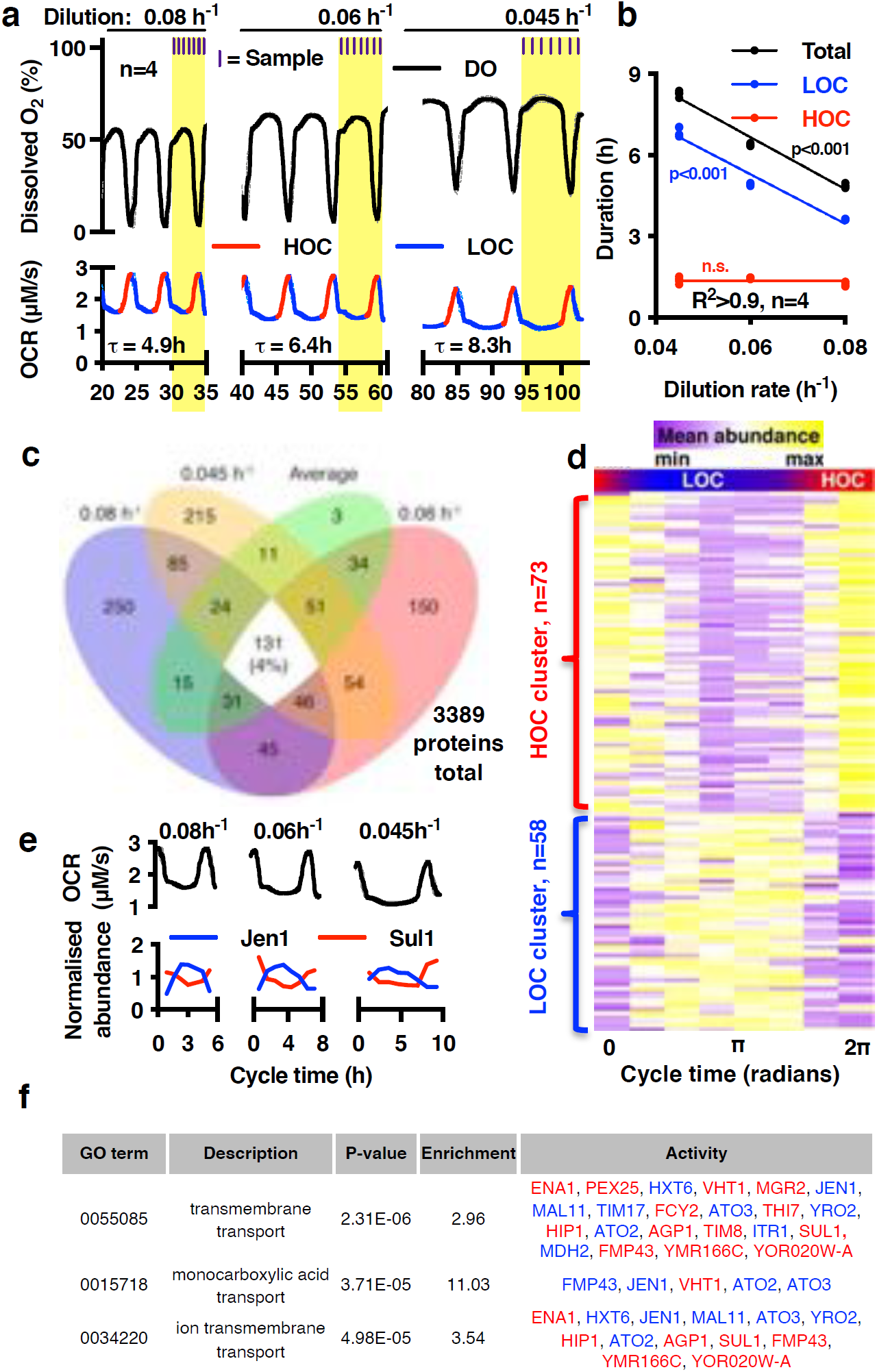
Rhythms of *S. cerevisiae* oxygen consumption and protein abundance at 3 dilution rates. **a,** The period (τ) of oscillation across high and low O_2_ consumption (HOC, LOC) varies with dilution rate (mean±SEM), sampling times indicated. **b**, the duration of LOC, not HOC, varies with dilution rate (p-value: straight line vs. horizontal line fit). **c,** Of 3389 proteins detected by quantitative mass spectrometry, only 4% were consistently rhythmic (varied by >33% across all conditions). **d**, Heatmap showing mean abundance of consistently rhythmic proteins clustered with either LOC or HOC. **e,** Oxygen consumption and mean-normalized protein abundance, for representative examples of HOC (high affinity sulphate permease, Sul1) and LOC (monocarboxylate transporter, Jen1) phased proteins. **f,** Table showing the most highly enriched gene ontology processes for consistently rhythmic proteins, proteins within each GO term are coloured by their YRO phase of expression (Red, HOC; Blue, LOC). Enriched proteins support the conclusion that membrane transport is important for YROs.

### Rhythmic regulation of ion transport, metabolic flux and cytosolic granules

Previous work suggested that YROs might organise through differentially phased gene expression in order to minimize the cost of expressing large genes^5,7,10,16^, with the expression of up to 60% of cellular mRNAs changing over the course of each cycle ^5,10,17^. To gain mechanistic insight, we measured protein, ion and metabolite content by mass spectrometry at multiple time points over the respiratory cycle, from 4 independent biological replicates at three dilution rates. Given the substantial proteomic coverage, we were surprised to find that only 4% of detected proteins varied with biologically significant amplitudes (>33%^18^) across all three dilution rates. This argues against a central role for dynamic changes in gene expression across the YRO, as does the poor correlation between the rhythmic amplitude, stability, abundance, size of each protein, or the energetic cost associated with production (Fig.1c-e, Extended Data Fig. 2,3). Of the few proteins whose abundance was rhythmic at the three dilution rates, unbiased k-means analysis suggested two clusters corresponding directly with LOC and HOC, with gene ontology analysis revealing a significant enrichment for terms associated with transmembrane transport of ions and carboxylic acids (Fig. 1f, Extended Data Fig. 2d). For example, the abundance of the sulphate transporter Sul1 clusters with HOC, whereas the monocarboxylate/proton symporter Jen1 clusters with LOC (Fig. 1e).

Our proteomic analyses suggested an unexplored requirement for differential ion transport during the YRO. Consistent with this, elemental and metabolite analysis revealed striking >2-fold variations in cellular osmolytes (K^+^, betaine, choline) over the oscillation, which accumulated during LOC and decreased during HOC (Fig. 2a, Extended Data Fig. 4,5). Increased osmolyte export was also detectable in the extracellular media, resulting in transient spikes in osmolality during HOC (Extended Data Fig. 5c).

**Figure 2.**
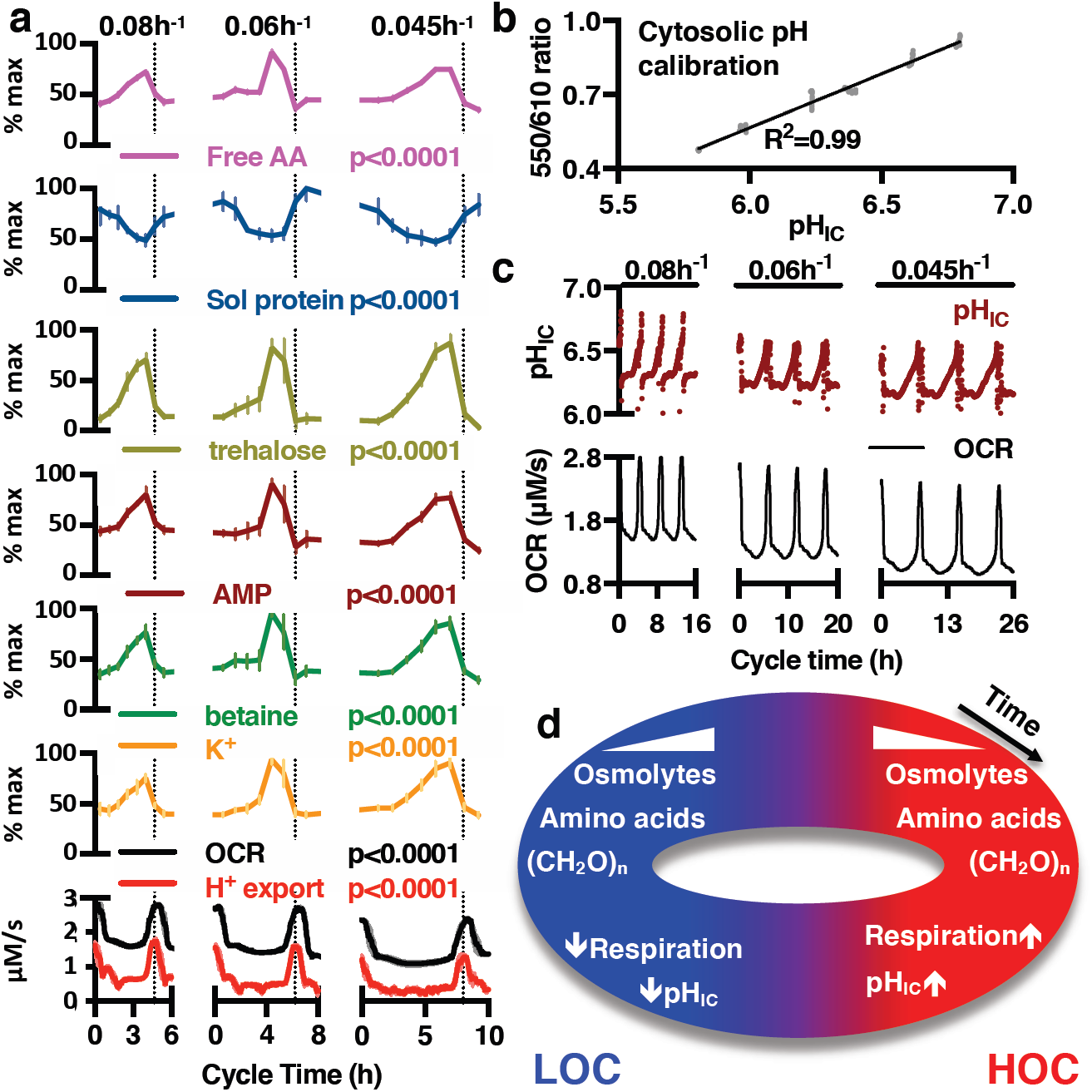
Consistent variation of critical metabolites, transport processes and soluble protein across the YRO. **a,** There are consistent phase relationships are between intracellular free amino acids, soluble protein, trehalose (storage carbohydrate), AMP, betaine, K^+^, OCR and H^+^ export under all conditions (mean±SEM, n=4, 2-way ANOVA_Time_ p-value shown). **b**, Calibration curve for firefly luciferase emission ratiometric reporter of cytosolic pH. **c,** Intracellular pH oscillates as a function of YRO phase under all conditions (representative data). **d**, Summary of key events that occur during HOC and LOC.

**Figure 3.**
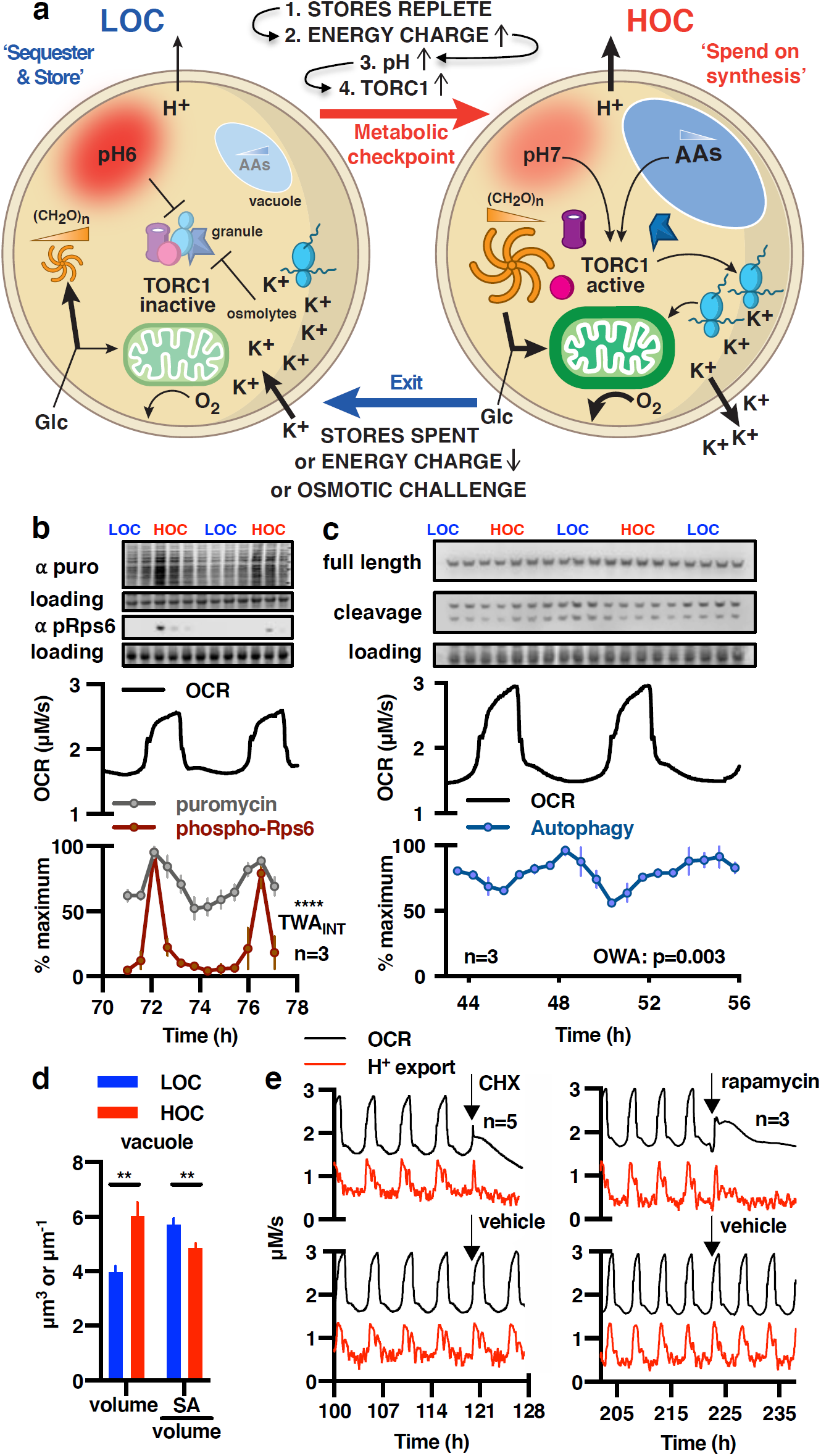
A pH-dependent, TORC1-dependent metabolic checkpoint switches between protein synthesis in HOC and autophagy in LOC. **a,** Model: cells accumulate carbohydrates (CH_2_O)_n_, amino acids and osmolytes during LOC and consume/export them in HOC to sustain translational bursts and osmostasis. Replete stores increase H^+^ export, a pH-dependent checkpoint activating TORC1 and releasing BMC proteins. HOC ends when stores are exhausted, see Extended Data Figure 7 for more detail. **b,** Puromycin incorporation assay and immunoblot for TORC1 activation (phospho-Rps6, Ser235/236) reveal translational bursting in HOC (mean±SEM, TWA_INT_: two-way ANOVA_INTERACTION_), total protein loading control. **c,** Immunoblots for cleaved/full-length Pgk1-GFP reveal increased autophagy during LOC (mean±SEM, OWA: one-way ANOVA). **d**, Differential variation in vacuole volume vs. surface area:volume ratio, mean±SEM, unpaired t-test, N=7 for HOC and LOC, n>68 cells/image. **e**, Acute inhibition of protein synthesis (CHX, 25µg/mL cycloheximide) or TORC1 activity 200nM rapamycin) during HOC immediately terminates HOC and abolishes the YRO, representative OCR and H^+^-export traces are shown.

Most active transport in yeast occurs through proton-coupled secondary active transport, driven by a difference of 3 pH units across the plasma membrane, that functions similarly to Na^+^ in mammalian cells ^19^. This ∼1000-fold gradient is generated by the essential ATP-dependent H^+^-pump, Pma1, which constitutes 15-20% of all yeast membrane protein ^20^ and consumes 20-40% of cellular ATP ^21^. The large variation in cellular osmolyte content over the YRO strongly suggested differential rates of H^+^ export. To test this, we derived the H^+^-export rate from the volume of NaOH required to maintain constant extracellular pH, as a proxy for Pma1 activity. Our data revealed a robust rise in cellular H^+^-export during HOC, which increased >2-fold in parallel with oxygen consumption, within each cycle (Fig. 2a, Extended Data Fig. 4a,b). Cytosolic pH functions as a second messenger in yeast ^22^, so we investigated whether intracellular pH mirrored the observed changes in H^+^-export. To do this we developed a ratiometric luciferase-based assay for real-time measurement of cytosolic pH over the YRO. We found that increased H^+^ export was accompanied by a significant rise in cytosolic pH (pH_cyto_) of >0.5 units (>3-fold drop in [H^+^]), irrespective of dilution rate (Fig. 2c), which rapidly returned to the lower pH at the HOC-to-LOC transition.

In agreement with previous work ^23^, >90% of other cellular metabolites we detected showed consistent variation across the YRO at each dilution rate (Extended Data Fig. 5), and unbiased k-means analysis again suggested two clusters, corresponding to HOC and LOC. Metabolites in the major cluster increased during LOC and fell during HOC (Extended Data Fig. 5d,e). Of particular note, amino acids and storage carbohydrates such as trehalose, and indicators of low energy charge such as AMP, all showed similar profiles to cellular osmolytes – increasing by 2-3-fold during LOC and then decreasing in HOC (Fig. 2a), whereas acetate and intermediates of phospholipid synthesis showed opposite profiles.

During glucose limitation or osmotic stress, yeast sequester proteins and mRNAs within non-membrane bound ribonucleoprotein biomolecular condensates (BMCs), including stress granules and p-bodies, whose formation is stimulated by pH_cyto_, and glucose starvation, and maintained by high osmotic potential ^24,25^. We predicted that the dramatic decrease in cellular osmolyte content and increase in cytosolic pH during HOC would increase the fraction of ‘soluble protein’ liberated from BMCs. We observed >2-fold variation in soluble cellular protein content, coinciding with HOC, whereas total cellular protein and protein in the media showed no significant variation (Fig. 2a, Extended Data Fig. 6a,b). Gene ontology analysis of soluble proteins with greatest variation between HOC and LOC revealed significant enrichment for protein chaperones, proteasomes, ribosomal subunits and key metabolic enzymes such as enolase (Eno1/2), pyruvate kinase (Cdc19) and decarboxylase (Pdc1), (Extended Data Fig. 6c). These proteins have previously been found in BMCs ^25-27^

### A testable model of the YRO

To explain our observations, we considered that protein synthesis is the most energetically expensive process that cells undertake, requiring high ATP turnover and tRNAs charged with amino acids^28^. The assembly of macromolecular protein complexes (e.g., ribosomes, proteasomes, electron transport complexes) is particularly challenging since subunits must be expressed stoichiometrically, at the same time, and without exceeding chaperone capacity, otherwise they are wastefully degraded or misfold and aggregate, with associated fitness costs^28-30^. Protein synthesis, folding and complex assembly are also sensitive to molecular crowding and osmolyte concentration^25,31^. Indeed, hyperosmotic challenge leads to degradation or sequestration of macromolecular solutes and export of osmolytes to maintain osmotic potential^32,33^. Since macromolecular complexes account for ∼40% of all cellular protein ^34,35^, efficient complex assembly is particularly challenging for yeast under nutrient-poor conditions, when cells carry out autophagy to generate free amino acids that serve as metabolic substrates for ATP production and accumulate carbohydrate stores of glycogen and trehalose. These stores are rapidly mobilised to fuel anabolic metabolism and cell cycle progression upon the return to growth^36,37^.

Target-of-Rapamycin Complex 1 (TORC1) is the master regulator of protein homeostasis that controls the switch between anabolic protein synthesis and catabolic autophagy ^38-40^. In yeast, TORC1 activity is regulated by glucose via increased pH/Gtr1-signalling, and indirectly by amino acid availability and energy charge via Gcn2 and Snf1/AMPK, respectively ^22,27,40,41^. TORC1 activity is also sensitive to molecular crowding, pH and osmolality ^31,39,42^. Coincidence detection through TORC1 ensures that the high translation rates required for efficient protein complex biogenesis only occur when sufficient energetic and biosynthetic resources are available.

In light of these well-established features of yeast cell biology, our observations suggest a mechanistic basis for understanding YROs, where oscillations ultimately arise from the selective pressure for efficient protein synthesis when nutrients are limiting (Fig. 3a, Extended Data Fig. 7). In this model, LOC is the default ‘sequester and store’ state, in which low amino acid availability, energy charge and pH_cyto_ inactivate TORC1, thus facilitating ATP generation through respiration of acetate and autophagic products while protein synthesis is low ^22^. Low pH_cyto_ also promotes progressive sequestration of macromolecules and glycolytic enzymes, such as Cdc19, into BMCs ^27^ and cytosolic removal of these colloidal macromolecular solutes stimulates osmolyte accumulation to maintain osmotic homeostasis. Respiration of acetate, autophagic and residual glycolytic products generate sufficient ATP to fuel basal Pma1-mediated H^+^ export, but reduced activity of sequestered glycolytic enzymes redirects the bulk of available glucose towards polysaccharide, lipid and nucleotide biosynthesis. The latter two are favoured because more glucose is available for NADPH and ribose production *via* the pentose phosphate pathway, increasing production of the essential building blocks for cell growth during G_1/2_ and S-phase of the cell cycle, respectively.

When carbohydrate stores are replete, surplus glucose and ATP stimulates increased H^+^ export by Pma1 ^43^, thereby raising pH_cyto_. Increased pH_cyto_ triggers the checkpoint for entry into HOC by stimulating a feed-forward switch that liberates glycolytic enzymes such as Cdc19 from BMCs to increase glycolysis, increasing energy charge and further elevating pH_cyto_. Increased pH_cyto_ also stimulates TORC1 to license increased protein synthesis, which is sustained because high amino acid levels and ATP:ADP/AMP ratio relieve TORC1 inhibition *via* Gcn2 and Snf1/AMPK pathways. Increased translation gradually consumes stored vacuolar amino acids, while energetic requirements are met by liquidation of trehalose and glycogen stores that fuel increased glycolysis and respiration. Liberation of chaperones, proteasomes and other quality control factors from BMCs aids efficient protein synthesis, assembly of protein complexes and turnover of damaged or misfolded proteins. To maintain osmotic homeostasis, the large increase in osmotic potential that would result from increased cytosolic macromolecular solutes during HOC is buffered by export of osmolytes (K^+^, betaine, choline) to maintain the cytosolic activity of water. In this model, rhythms of respiration and metabolism are ultimately driven by the bioenergetic demands of increased TORC1-stimulated protein synthesis, triggered by elevated pH_cyto_. DNA replication and lipid synthesis are restricted during HOC (Extended Data Figs.1 & 5), because cellular resources are directed towards protein production.

According to our model (Extended Data Fig. 7), the end of HOC occurs due to 1) cytosolic acidification due to insufficient ATP (a result of depleted carbohydrate stores and reduced glucose supply), 2) insufficient O_2_ supply which also reduces ATP production and/or 3) exhaustion of amino acid or osmolyte stores, resulting in Gcn2 activation or osmotic challenge, respectively. The period of oscillation is therefore determined by the amount of time taken to replenish osmolyte, amino acid and carbohydrate stores during LOC, and the (normally) invariant duration of HOC reflects a consistent time taken to ‘spend’ stored carbohydrates, osmolytes and/or amino acids on protein synthesis. We would therefore expect that exit from HOC will be brought forward by acute hyperosmotic stress, translational or respiratory inhibition.

### YRO model validation

This model is consistent with available data and makes many testable predictions. For example, we observed significantly higher rates of protein synthesis and TORC1 signalling during HOC, whereas autophagy was more active during LOC (Fig. 3b,c). Autophagic breakdown products such as free amino acids are stored in the cell vacuole^44^, and we observed significant differences in vacuolar morphology between LOC and HOC corresponding with differential autophagic flux across the YRO (Fig. 3d).

To test our prediction that the energetic demands of increased protein synthesis drive the characteristic increase in oxygen consumption during HOC, we added cycloheximide (CHX) or rapamycin to cells, to inhibit translation or TORC1 activity, respectively. Confirming expectation, both drugs immediately curtailed the anticipated increase in oxygen consumption and proton export during HOC (Fig. 3e), and abolished subsequent oscillations, but without similar effect on basal levels of respiration. At lower, non-saturating concentrations, rapamycin addition during HOC shortened the period of YRO oscillations, but critically the first effect of drug treatment was observed during the next HOC rather than the intervening LOC (Extended Data Fig. 8a,b). By our model LOC is expected to be insensitive to acute TORC inhibition but reduced in length after a truncated HOC, since stores will not be fully depleted and so take less time to replenish. These observations strongly support our hypothesis that TORC1 inactivation suppresses protein synthesis during LOC, and that TORC1 activation triggers the increase in translation that drives the demand for greater respiration during HOC.

We next sought to assess the extent of differential BMC sequestration and glycogen storage over the YRO. Consistent with expectation, the stress granule marker, Pab1, was significantly more diffuse during HOC, whereas brighter foci were evident during LOC (Fig. 4a). Moreover, the amount of cellular glycogen fell by 40% during HOC (Fig. 4b), with a profile very similar to that of trehalose (Fig. 2a), the other major storage carbohydrate.

**Figure 4.**
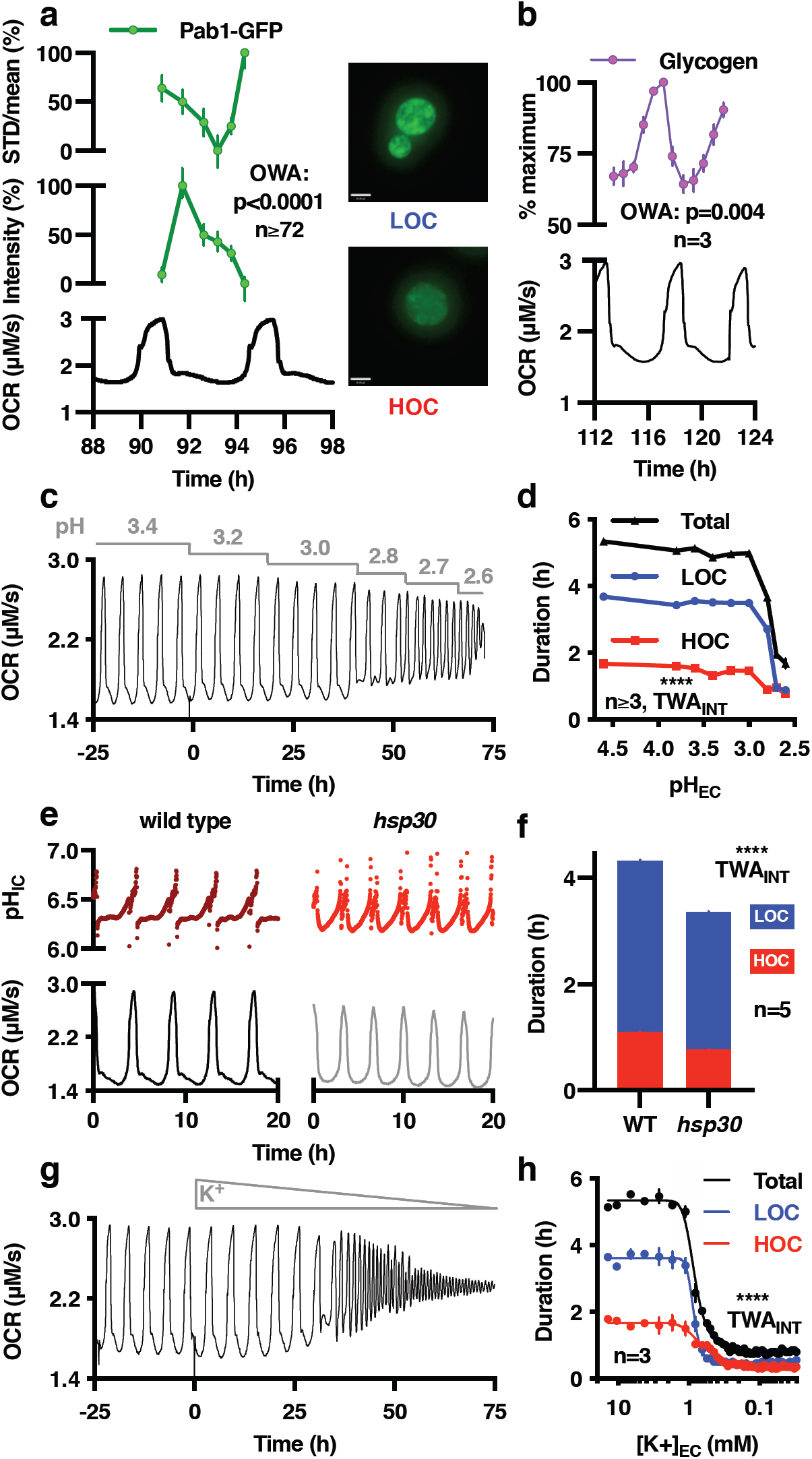
YROs differentially regulate stress granule formation and glycogen storage, and are extremely sensitive to perturbation of H^+^ and K^+^ transport. **a,** The intensity and distribution (STD/mean) of stress granule marker Pab1 (Pab1-GFP signal) varies over the YRO, with more foci during LOC and more diffuse during HOC, supporting dynamic variation in stress granule formation (mean±SEM). **b**, Cellular glycogen stores increase during LOC and decrease during HOC (mean±SEM). Liquidation of storage carbohydrates is likely to fuel translational bursting during HOC. **c**,**d** Decreasing extracellular pH reduces the period of the YRO duration (representative OCR, mean±SEM quantification). **e,f**, *HSP30* mutants have truncated oscillations (representative OCR, mean±SEM quantification). **g,h,** Extracellular K^+^ concentration determines YRO period duration (representative OCR, mean±SEM). This is unlikely to be due to loss of cell viability as YROs are rapidly restored when potassium becomes available (representative OCR, mean±SEM quantification).

Given the pivotal role of Pma1-mediated H^+^ transport in our model, we tested a prediction that the period of oscillation should be sensitive to increases in pH gradient across the plasma membrane because more ATP turnover will be required to maintain pH_cyto_, in both HOC and LOC. Concordantly, we found YROs run >2-fold faster with decreasing, but not increasing, extracellular pH (Fig. 4c,d). Our model also predicts increased Pma1 activity will elicit shorter cycles, as this accelerates the HOC to LOC transition. We tested this with mutants of *hsp30*, a negative regulator of Pma1^45^, observing shorter, truncated cycles of oxygen consumption and pH_cyto_ (Fig. 4e,f).

From the model, dynamic import and export of osmolytes during LOC and HOC, respectively, is critical for buffering cytosolic osmotic potential: permitting cycles of TORC1 activity and reversible macromolecular sequestration in BMCs/stress granules, both of which are critical for HOC translational bursting and associated respiration increase. Consistent with this, an acute osmotic challenge during HOC, which both inhibits TORC1 activity^31^ and opposes protein liberation from BMCs, resulted in immediate exit from HOC (Extended Data Fig. 8c).

To functionally validate the importance of intracellular osmolyte accumulation to YRO period determination, at the transitions between LOC and HOC, we manipulated the major intracellular osmolyte K^+^. The model predicts that decreased osmotic buffering will shorten the duration of both HOC and LOC. This is because insufficient K^+^ accumulation in LOC means that HOC cannot be sustained, resulting in early osmotic challenge, whereas LOC will be shorter because stored carbohydrates/amino acids were not exhausted during the previous HOC. This in turn, will reduce the time taken to reach the pH_cyto_ threshold for HOC entry in the next cycle. To test this, the infeed was switched to media where K^+^ was replaced with Na^+^. As extracellular K^+^ decreased, we indeed observed a dramatic and reversible shortening of YRO period; whereas depleting Mg^2+^, another essential metal ion but without significant osmotic function, simply abolished oscillations (Fig. 4f,g, Extended Data Fig. 8d,e). Conversely, pulse addition of K^+^ increased YRO period, as well as basal OCR, HOC oxygen consumption and duration, because increased K^+^ availability facilitates greater osmolyte accumulation during LOC allowing HOC to be sustained for longer (Extended Data Fig. 8f,g).

### Physiological consequences

We next explored the physiological consequences of our YRO model, which predicts that the yeast should be more sensitive to a heat stress during HOC, as increased cytosolic protein concentration, reduced osmotic buffering capacity and lower trehalose abundance should render cells more susceptible to protein denaturation. To test this, cells sampled from different points of the YRO were immediately subjected to an acute heat shock, and survival measured under standard growth conditions. Consistent with prediction, a two-fold difference in viability was observed between cells harvested at the minimum and maximum oxygen consumption rate (Fig. 5a). We then tested whether translational bursting during HOC is sensitive to acute perturbation of the transmembrane pH gradient and to osmotic challenge, as would be expected. For this assay, we measured protein synthesis by puromycin incorporation. Cells were removed from the bioreactor during HOC and transferred immediately into media of different pH or osmolality. Consistent with expectations, higher pH acutely increased translation whereas lower extracellular pH and hyperosmotic media reduced it (Fig. 5b, Extended Data Fig 9).

**Figure 5.**
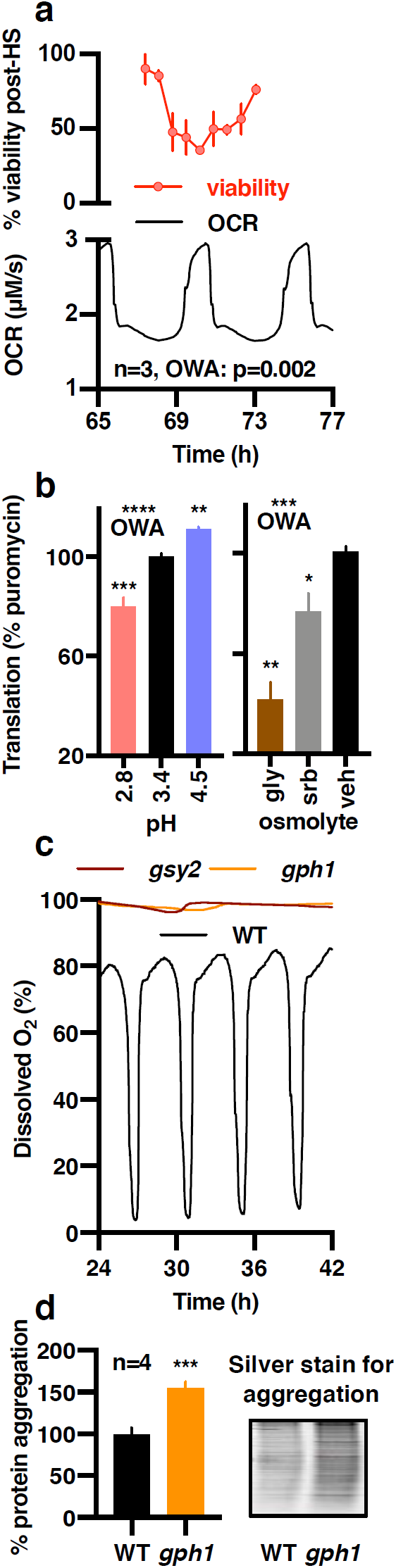
The YRO regulates resistance to heat stress and protein homeostasis. **a,** Viability of cells removed from the bioreactor after heat treatment (55°C, 2 mins) is greatest at the end of LOC, when the abundance of trehalose and osmolytes are greatest. Percentage of heat-treated cells, corrected for viability of non-heat treated cells harvested at the same time. **b**, Sensitivity of HOC protein synthesis rate to pH and hyperosmotic stress assayed by puromycin incorporation (gly, 10% glycerol; srb, 1 M sorbitol, n=4). **c**, Strains deficient in glycogen synthesis (*gsy2*) or glycogen breakdown (*gph1*) do not initiate YROs and, **d**, *gph1* strains accumulate aggregated protein, showing that glycogen breakdown is necessary for proteostasis. Representative silver-stained gel (mean±SEM, unpaired t-test).

Moving beyond the mechanistic determinants and ramifications of this biological rhythm, our model makes an explicit prediction about its functional utility: translational bursting requires high ATP turnover that can only be sustained by the rapid mobilization of glucose from carbohydrate stores. Therefore, glycogen metabolism mutants will have no temporal organization to the synthesis of proteins or respiration. This will render nascent proteins more likely to exceed chaperone capacity, and/or fail to find a binding partner, and so increase misfolded and aggregated proteins. Consistent with this, glycogen storage or consumption mutants did not exhibit respiratory oscillations and showed a 50% increase in protein aggregation (Fig. 5c,d), indicating a profound deficiency in protein homeostasis^29,30^.

## Conclusion

We provide an experimentally-derived, predictive and testable model of the yeast respiratory oscillation wherein central metabolism, signal transduction, active transport, and macromolecular condensation are temporally organized to accommodate the bioenergetic demands of protein synthesis. This model accounts for the period of oscillation in that HOC duration depends on the amount of time taken to satisfy one or more of the conditions for HOC exit, whereas the duration of LOC depends on the time taken to accumulate carbohydrate.

Eukaryotes have evolved under conditions where nutrients are limiting and environmental conditions vary. Given the high energetic cost of macromolecular complex assembly, and the selective pressure for efficient use of resources, we propose that oscillations in global protein synthesis rate confer a general fitness advantage, realized in many other biological contexts, that is supported by the same ‘save and spend’ partitioning of metabolic resources we have observed in yeast.

Aspects of our model that involve H^+^ gradients and vacuoles are unlikely to apply to eukaryotic cell types that lack cell walls or use Na^+^ instead of H^+^ gradients. However, we speculate that dynamic transport of cellular osmolytes to buffer osmotic homeostasis against changes in macromolecular condensation, and dynamic rerouting of metabolic flux to increase ATP production during TORC1-dependent translational bursting, are essential for the temporal organization of proteome remodeling, and therefore common to most eukaryotic cells. Cell-autonomous rhythms of TORC activity, protein synthesis, potassium transport and cellular respiration have been all been observed in mammalian cells over the course of the circadian cycle ^46-55^. We therefore speculate that the central utility of biological oscillations such as the YRO, and circadian rhythms throughout the eukaryotic lineage, is to facilitate the efficient utilisation of metabolic resources in order to minimize the cost of protein homeostasis.

## Acknowledgements

We thank Siu-Hong Ho (CCTI Flow Core), Marcus Mercado (Eppendorf), Gerben van Ooijen, Liz Miller and David Tourigny for useful discussions, members of the Pon Laboratory (particularly Theresa Swayne and Jie-Ning Yang) and Columbia University Confocal and Specialised Microscopy Shared Resource for assistance with imaging and image analysis and members of the O’Neill and Causton labs for their patience. The CEN.PK113-7D strain was a gift from Peter Kötter (Johann Wolfgang Goethe University, Frankfurt am Main). JBR is supported by the Center for Molecular Biosciences, MTSU. CF and JSO are supported by the Medical Research Council (MRC_MC_UU_12022/6 & MC_UP_1201/4, respectively).

## Author contributions

J.S.O. and H.C.C. conceived the idea, performed experiments and analysis, and wrote the manuscript. N.P.H., J.B.R., R.S.E., A.D.B., S.C., J.D., A.S.H.C. & C.F. performed experiments and analysis, and made valuable intellectual contributions.

