## Supplemental Data for "Eukaryotic cell biology is temporally coordinated to support the energetic demands of protein homeostasis"

**Figure Legends**

**Extended Data Figure 1, Relating to Figure 1. The YRO comprises phases of higher and lower oxygen consumption and gates S-phase entry but not mitosis, and is therefore distinct from the cell division cycle**

**a,** Individual dissolved oxygen traces from the four bioreactor vessels used for harvesting of cells for multi-omic analysis. **b,** Reoxygenation of media (without cells) over time under conditions used for YROs. **c**, Dissolved $O_2$ quickly equilibrates with that in the gaseous phase, so under constant aeration the rate at which dissolved $O_2$ increases, decreases as $[O_2]$ approaches saturation (n=4). **d**, As $O_2$ is supplied at a constant rate, the rate at which $O_2$ is consumed ($O_2$ consumption rate, OCR) to achieve any given steady state $[O_2]$ can be interpolated from the standard curve. **e**, To allow comparison between experiments, we define HOC as the time when the rate of change in oxygen consumption rate is above zero, LOC when below zero. HOC and LOC durations may readily be quantified from the first derivative of OCR (mean±SEM, for the 4 replicates shown in **a**). **f**, DNA replication is inhibited at the transition into and during HOC, resulting in an oscillation in the relative proportion of 1C to 2C cells ($G_1$:$G_2$). There is no significant variation in cell number across the YRO that is consistent for all three dilution rates, however. This means that any variation observed across the YRO at all three dilution rates cannot be attributed simply to synchronized mitosis or variation in cell number (mean±SEM, n=4, 2-way ANOVA for time effect reported). **g**, Representative flow cytometry data showing the intensity of the propidium iodide signal (DNA) vs. forward scatter for populations of yeast oscillating at 0.08 dilutions/h.

**Extended Data Figure 2, Relating to Figure 1. Quantitative proteomics reveal modest changes in protein abundance across the YRO with clusters that correspond to HOC and LOC.**

**a**, Plot of fold change vs. detected proteins for three dilution rates. Informed by quantitative genome-wide measurements of the intrinsic noise of gene expression in yeast[18], and given that our approach will not detect very low copy number proteins, we chose a conservative threshold for biological significance of 1.33. This means that any protein whose abundance does not change by >33% over the YRO at all three dilution rates, as well as the average abundance across all dilutions, is not considered to be consistently rhythmic. **b**, Amongst the consistently rhythmic proteins, unbiased k means cluster analysis provided no strong support for any specific number of clusters between 2-10, as revealed by the 'break point' in a plot of inter-cluster variation versus cluster number. We therefore selected the simplest model, *i.e.*, two clusters. The temporal profiles of proteins in these two clusters correspond with the relative phases of HOC and LOC. **c**, Plot of normalized protein abundances at each dilution rate vs. OCR (top, repeated from Figure 1a), stratified by profile. Three randomly selected examples highlight proteins whose expression peaks during LOC-or HOC. Note that the majority of detected proteins 2247/3389 varied by <33% at all 3 dilution rates, whereas proteins that were consistently rhythmic typically showed modest changes in abundance. d, The most enriched non-redundant Gene Ontology processes for consistently rhythmic proteins in the HOC and LOC clusters reveals differential regulation of transporters in both YRO phases. Also see Supplementary Data Table 1.

**Extended Data Figure 3, Relating to Figure 1. Proteomics analysis reveal weak correlations between fold change in protein abundance across the YRO with their stability, abundance, size or cost.**

**a-d,** The expression of most mRNAs varies over the YRO [5,10], yet the abundance of most proteins does not, implying that transcriptional rhythms may function to maintain steady state protein levels, rather than to drive rhythms in the activity of the encoded protein. Consistently rhythmic proteins were enriched for transport processes which implies that differential transporter activity over the YRO is relevant to its mechanism and biological function. However, it was equally plausible that more rhythmic proteins were simply those with shorter half-life, or those with higher copy number, such that changes in abundance would be more readily detectable. Because larger proteins are more costly to produce [17] another possibility is that natural selection has acted to suppress the production of large proteins at stages of the YRO when they are not needed, leading to rhythms in abundance. Considering all 3389 detected proteins, however, poor correlations were observed between each protein's relative amplitude and its half-life, abundance, size or the energetic cost of its synthesis (size x abundance). Data combined across all three dilution rates (mean±SEM, n=3). Since all 4 simpler hypotheses were rejected by these analyses, we consider that the over-representation of transporters among consistently rhythmic proteins occurs in response to the requirement for differential regulation of transporter activity over the YRO. **e,** Heatmap showing all consistently rhythmic proteins without clustering, normalized to the relative change for each protein over the course of the oscillation.

**Extended Data Figure 4, Relating to Figure 2. $H^+$ and $K^+$ export vary consistently over the YRO**

**a,** NaOH is pumped into bioreactor vessels to maintain the media at pH 3.4; greater NaOH addition occurs during HOC compared with LOC (mean±SEM, n=4). **b,** From the rate of NaOH addition and its concentration, the rate at which the cell population exports $H^+$ is readily interpolated. $H^+$ export rate changes in parallel with the oxygen consumption rate (OCR) (mean±SEM, n=4). **c,** and **d,** Samples of cells undergoing YROs were harvested across four replicates at three dilution rates, as described, and subject to analysis of intracellular metal ion content by ICP-MS using $^{43}Ca$, $^{44}Ca$, $^{59}Co$, $^{63}Cu$, $^{65}Cu$, $^{57}Fe$, $^{39}K$, $^{24}Mg$, $^{55}Mn$, $^{60}Ni$, $^{31}P$, and $^{66}Zn$. Only the intracellular concentration of potassium ions shows variation significant for time but not dilution or interaction effect (mean±SEM, n=4, 2-way ANOVA reported). N.B. Due to the preparative method used for these yeast cell samples, $Na^+$ and $Cl^-$ could not be measured (which was required to make them compatible with the metabolomic and proteomic data).

**Extended Data Figure 5, Relating to Figure 2. The majority of detected cellular metabolites vary consistently over the YRO**.

**a,** The abundance of most detected metabolites significantly and consistently varies across the YRO (mean±SEM, n=4, 2-way ANOVA reported). 82/89 of the metabolites identified in each replicate at every time point showed a time effect by two-way ANOVA with $p<0.05$, we describe these as consistently rhythmic N.B. unlike the other metabolites, acetate was not validated against external standards but its profile is entirely consistent with previous reports[5]. **b**, Cellular ATP content varies over the YRO, measured enzymatically as previously[46] (mean±SEM, n=4, 2-way ANOVA reported). **c**, The osmolality of the extracellular media transiently increases during HOC (mean±SEM, n=4, 2-way ANOVA reported). **d**, As with consistently rhythmic protein profiles (Extended Data Fig. 2b), unbiased k means cluster analysis of consistently rhythmic metabolites suggests that two clusters of temporal profile were appropriate, as revealed by the 'break point' in a plot of inter-cluster variation versus cluster number. **e,** Heat map for consistently rhythmic metabolites associated with Cluster 1 or 2, normalized to the average minimum and maximum detected abundance of each metabolite for each YRO cycle. Cluster 2 suggests an association of phospholipid synthesis with LOC, whereas profiles of metabolites in cluster 1 are similar to $K^+$, peaking around the LOC=>HOC transition and falling as OCR increases. This includes amino acids, organic osmolytes and storage carbohydrates.

**Extended Data Figure 6, Relating to Figure 2. Release of protein from cytosolic storage/stress granules, or BMCs, varies consistently over the YRO.**

**a,** The amount of protein resolubilized in reducing 8 M urea buffer following hot ethanol precipitation (soluble protein), from the equal numbers of cells, changes by more than 2-fold across the YRO, while the concentration of total cellular protein or protein in the media does not change significantly (mean±SEM, n=4, 2-way ANOVA time effect p-value reported). Precipitation in hot ethanol favours intramolecular disulphide formation for hydrated 'soluble proteins' in the cytosolic compartment, compared with an increased relative likelihood of intermolecular disulphide formation for proteins in BMCs. Upon reconstitution in reducing 8 M urea buffer at 37°C, most BMC disulphides are less solvent-accessible to the reductant (10 mM TCEP), and more likely to remain within insoluble aggregates that are removed by centrifugation, whereas precipitated proteins with accessible disulphides that were not within proteinacious non-membrane bound are more likely to go into solution and therefore remain in the supernatant upon centrifugation[56]. **b**, Representative coomassie-stained gel showing soluble protein extracted from equal numbers of cells harvested across YRO cycles at three dilution rates (n=4). **c**, Mass-spectrometry of proteins with highest variation across the YRO (*labelled 1 to 4) reveals enrichment for gene ontology terms associated with protein folding and degradation, as well as catalytic and oxido-reductase activity. From the proteins that were identified in each band, the most enriched GO term (function) is reported, with corrected p-value and identified proteins. 4* Ribosomal proteins were identified in band 4, however, they are not significantly enriched, due to the large number of ribosome-associated proteins in cells at all times.

**Extended Figure 7, Relating to Figure 3. A detailed, testable and experimentally-derived YRO model**

Green arrows/lines represent activation/repression, red arrows represent ATP production/stimulation of ATP production, Black arrows represent predicted metabolic flux, see key for further details.

**Extended Figure 8, Relating to Figures 3 and 4. Further experimental perturbation of the YRO.**

**a,b,** Inhibition of TORC1 activity during HOC by addition of rapamycin transiently decreases the period and amplitude of oscillation. YROs recover as the drug is diluted out of the reactor vessel. The duration of LOC is affected more than HOC, likely because the resource required to support translation is not exhausted during HOC and is therefore replenished faster. Critically, the first LOC after rapamycin addition is not affected, whereas the next HOC is, consistent with differential regulation of TORC1 activity. **c,** Osmotic stress promotes premature exit from HOC, likely due to increased macromolecular crowding, which inhibits TORC1 and opposes the liberation of proteins from BMCs. **d,e,** The effect of potassium depletion is ion-specific and is not due to loss of viability, as a return to standard inflow media (containing 14mM $K^+$) rapidly restores normal oscillations. **f**, **g,** The addition of KCl to cells undergoing HOC increases the duration of HOC, OCR and basal respiration rate, this is the opposite of the response to potassium depletion.

**Extended Figure 9, Relating to Figures 3 and 5. Relationships between vacuole morphology and protein homeostasis with the YRO, extracellular osmolality and pH.**

**a, b** Full immunoblots shown in Figure 3b and c. Red lines indicate the regions shown. **c,** Cells sampled from the bioreactor during LOC and HOC have vacuoles of different shapes (vacuoles stained with CMAC, representative of images quantified in Figure 3d). **d,** Strains deficient in the breakdown of glycogen (*gph1*) harbour more aggregated protein than wild type strains. Silver stained gel, quantified in Figure 5d, showing data from 4 biological replicates. **e,** Cells sampled from the bioreactor in mid-HOC incorporate less puromycin when diluted into media containing 10% glycerol or 1 M sorbitol, while cells sampled during early HOC incorporate less puromycin when diluted into media of pH 2.8 and more puromycin when diluted into media of pH 4.8. Data quantified in Fig. 5b.

#### Methods

##### Bioreactor continuous measurement and discrete sampling

Respiratory oscillations were initiated and maintained as described [2,5] in a DASgip bioreactor. For generation of samples for high-throughput analysis (metabolomics, proteomics, ionomics and flow cytometry) the bioreactor was operated with 4 replicates in parallel at 220 sL/h aeration, 550rpm agitation, 1.5L media/vessel, at 0.08, 0.06 or 0.045 dilutions/h, 30°C, pH 3.4 was maintained by addition of 2M NaOH. Media is as described previously [45]. The dissolved oxygen ($DO_2$) traces are shown in Extended Data Fig. 1a. pH was monitored using a calibrated pH electrode and the pH maintained by automated addition of 2N NaOH. In 1.5 L media, 1 mL/h 2N NaOH neutralises $H^+$ production equivalent to 0.37 µM/s.

Dissolved $O_2$ was monitored with an $O_2$ electrode. To establish the relationship between relative $DO_2$ and oxygen consumption, the $O_2$ electrode was calibrated by allowing anoxic media to equilibrate with atmospheric $O_2$ under the conditions described above (100% = 210 µM $O_2$ at 30°C, Extended Data Fig. 1b). This permits oxygen consumption rate (OCR, µM/s) to be derived for any steady state concentration of $O_2$ (Extended Data Fig. 1d) where $O_2$ is removed from the system. To allow quantitative comparison between experiments, we defined HOC as a sustained increase in OCR such that the 10 min moving average of the first derivative of OCR (dOCR/dt) is >0 for ≥10 min and increases by >10% overall within each bout. The phase of LOC occurs between bouts of HOC.

At stated time points, 1.5mL samples were removed from the bioreactor and harvested by centrifugation (2840 g, 4°C, 3 minutes). The supernatant and pellet were flash frozen separately in liquid nitrogen and stored at -80°C. Cell pellets (60 µL volume) were then thawed on ice, and washed twice in 1 mL ice cold, isosmotic buffer X (25 mM iodoacetamide, 25 mM NaF, 25 mM $NaN_3$, $H_2SO_4$ added to pH 3.4, made up to 150 mOsm with 1 M sorbitol) to remove extracellular ions, proteins and metabolites and inactivate cellular enzymes. Pellets were resuspended in 1 mL buffer X, split into 3 aliquots, centrifuged, supernatant removed, and stored at -80°C for analysis by mass spectrometry.

Unless stated, all other bioreactor experiments took place under identical conditions except that 175 sL/h aeration and 1.0 L media/vessel were used, with appropriate $O_2$/pH calibrations and calculations also being performed. Drugs were dissolved in DMSO (Sigma D2650) and added by pulse addition of drug or vehicle to the bioreactor vessel and infeed media. Rapamycin (Sigma R0395) 15 nM or 200 nM final concentration, cycloheximide 25 µg/mL (Sigma C7698).

**Proteomics**

***Sample preparation***

20µl pellets of yeast were lysed in 100 µL lysis buffer (6 M urea, 2 M thiourea, 20 mM HEPES pH 8, with protease and phosphatase inhibitors) with 50 µL of 0.5mm glass beads by agitation using a Bioruptor Genie (Scientific Industries) at 4 °C (3 x 60 seconds with 5 mins on ice between runs). Lysates were clarified by centrifugation at 21000 g for 5 mins and total protein concentrations were determined by Pierce BCA assay (Thermo) The concentration of each sample was normalised to 200 µg at 1 mg/mL. For each dilution rate, 100 µL of supernatant from 4 biological replicates pooled to generate samples representing 9 time points. A pooled cell extract reference sample was made by mixing 25 ul from each of the 9 samples.

***Protein digestion***

Samples in 200 µL lysis buffer were reduced with 5 mM DTT at 56 °C, 30 min and alkylated with 10 mM iodoacetamide in the dark at room temperature, 30 min. Samples were then digested with Lys-C (mass spectrometry grade, Promega), 133:1 (protein: Lys-C ratio, w/w) for 4.5h, 25 °C and then diluted from 8 M to 1.8 M urea with 20 mM Hepes (pH 8.5) and digested with trypsin (Promega) 100:1 (protein: trypsin ratio, w/w) overnight, 25°C. Digestion was stopped by the addition of trifluoroacetic acid (TFA) to 1% (final). Precipitates were removed by centrifugation (10000 rpm, 5min.). The supernatants were desalted using home-made C18 stage tips (3M Empore) containing 4 mg poros R3 (Applied Biosystems) resin. Bound peptides were eluted with 30-80% acetonitrile (MeCN) in 0.1% TFA and lyophilized.

***Tandem mass tag (TMT) labeling***

Peptide mixtures were re-suspended in 75 µL 3% MeCN and concentrations determined using the Pierce Quantitative Colorimetric Peptide assay (Thermo Scientific) according to the manufacturer's instructions, except the absorbance was measured using NanoDrop Spectrophotometers (Thermo Scientific) at 480nm. TMT 10plex reagent (Thermo Fisher Scientific) of 0.8 mg each was re-constituted in 41 µL anhydrous MeCN. 61.5 µL (1.5 x 0.8mg) of the reagent was used for each sample. The labeling reaction was performed in 150 mM triethylammonium bicarbonate, 1hr at room temperature and terminated by 15 min incubation with 9 µL 5% hydroxylamine. Labeled peptides were combined into a single sample and partially dried to remove acetonitrile in a SpeedVac. The labeled mixture was desalted using C18 stage tips, with 6.6 mg of R3.

***Proteome: off-line high pH reverse-phase peptides fractionation***

Approximately 100 µg of the labeled peptides were separated on an off-line, high-pressure liquid chromatography (HPLC). The experiment was carried out using XBridge BEH130 C18, 5 µm, 2.1 x 150mm (Waters) column with XBridge BEH C18 5 µm Van Guard cartridge, connected to an Ultimate 3000 Nano/Capillary LC System (Dionex). Peptides were separated with a gradient of 1-90% B (A: 5% MeCN/10 mM ammonium bicarbonate, pH8 [5:95]; B: 90% MeCN/10 mM ammonium bicarbonate, pH8, [9:1]) in 60 min at a flow rate of 250 µl/min. 60 fractions were collected, combined into 20 fractions and partially dried in a Speed Vac to about 50µL.

***LC-MSMS***
Liquid chromatography was performed on an Ultimate 3000 RSLC nano System (Thermo Scientific) fitted with a 100 µm x 2 cm PepMap100 C18 nano trap column and a 75 µm×25 cm reverse phase C18 nano-column (Aclaim PepMap, Thermo Scientific). Samples were separated using a binary gradient consisting of buffer A (2% MeCN, 0.1% formic acid) and buffer B (80% MeCN, 0.1% formic acid). Peptides were dissolved in solvent A and eluted with a step gradient of 5 to 50% B in 87-105 min, 50 to 90% B in 6-10 min, with a flow rate of 300 nL/min. The HPLC system was coupled to a Q-Exactive Plus mass spectrometer (Thermo Scientific) with a nanospray ion source. The mass spectrometer was operated in standard data dependent mode, performed MS full-scan at 350-1600 m/z range, resolution 140000. This was followed by MS2 acquisitions of the 15 most intense ions with a resolution of 35000 and NCE of 32%. MS target values of 3e6 and MS2 target values of 1e5 were used. Isolation window of precursor was set at 1.2 Da and dynamic exclusion of sequenced peptides enabled for 40s.

***Analysis***
The MSMS raw files were processed using Proteome Discoverer (v2.1, Thermo Scientific). MSMS spectra were searched against the reviewed *Saccharomyces cerevisae*, UniProt Fasta database (July 2017), using Mascot (version 2.4, Matrix Science) search engine. Carbamidomethylation of cysteines was set as fixed modification, while methionine oxidation, N-terminal acetylation (protein), phosphorylation (STY) and TMT6plex (peptide N-terminus and Lysine) as variable modifications. Other parameters were precursor mass tolerance, 10 ppm and fragment mass tolerance, 0.03 Da. Only peptides with FDR of 1% based on a target decoy approach (high confidence peptides) were included in the results. The output file from Proteome Discoverer, proteins table was filtered for proteins with FDR of 1% and exported as Excel files. The mass spectrometry proteomics data have been deposited to the ProteomeXchange Consortium via the PRIDE partner repository [57] (proteomecentral.proteomexchange.org) with the dataset identifier <PXD013653>.

Sample:pooled sample ratio(SPS) was calculated for each time-point and normalised so that the median SPS = 1. Sampling times were calculated from dissolved oxygen ($DO_2$) measurements and converted into radians where 2π radians is equal to the offset time where the correlation coefficient of a serial autocorrelation was at its maximum. Linear interpolation was used on each dataset to calculate the mean SPS across all dilution rates every 0.9 radians. The fold-change between the maximum and minimum SPS detected in the time course for each protein identified in all dilution rates was calculated (FC).

FC versus protein half-life and length in residues was calculated from [58]. FC versus relative protein abundance and protein cost (relative abundance x length in residues) was also calculated. Protein SPS profiles where FC>1.33 were clustered (k means, Hartigan Wong algorithm, R version 3.3.3). The between cluster sum of squares / total within cluster sum of squares was calculated for 1-10 clusters. Due to the lack of inflection points in the plotted data (Extended Data Fig. 2d) we used 2 clusters for our analysis where the greatest change in cluster/total sum sum-of-squares occurred. GO analysis was performed on each cluster independently, and when combined, using SGD GO Term Finder version 0.86 using the Process ontology with p<0.05 and default settings.

**Ionomics**

20 µL cell pellets were dissolved in 550 µL 65% HNO3, supplemented with 100 p.p.b. cerium, at 90 °C for 1 h then centrifuged at 18,000 g for 20 min to remove any debris, then diluted 1:12 in HPLC-grade water to give a final matrix concentration of 5% HNO3. Cellular elemental composition was determined by inductively-coupled plasma mass spectrometry (ICP-MS) on a Perkin Elmer Elan DRC II ICP-MS instrument in helium collision mode. Cerium was used to correct for dilution errors introduced during handling, then normalized to sodium in buffer X. The operator was blinded to the samples, which were randomized to avoid any effect of machine drift. Ion intensities were calibrated against 10x SPS-SW2 standard (LGC), which was injected 9 times during the run. The calibration was checked for accuracy with a second multi element standard purchased from SCP Science (Canada).

**Metabolomics**

***Metabolite extraction and analysis***

Cells were collected by centrifugation, the medium discarded and the samples extracted with 200 µl 80°C hot ethanol. Residual medium resulted in a final ethanol concentration of approximately 80%. The extract was heated for 2 min at 80°C, vigorously mixed on a vortex mixer and incubated for further 2 min at 80°C [59]. The extract was cleared of debris by centrifugation and stored at −80°C for subsequent analysis by liquid chromatography mass spectrometry (LC-MS).

LC-MS analysis was performed on a Dionex U3000 UHPLC system coupled to a Q Exactive mass spectrometer (Thermo Fisher Scientific). The liquid chromatography system was fitted with a SeQuant ZIC-pHILIC column (150 x 2.1 mm, 5 µm) and guard column (20 x 2.1mm, 5 µm) from Merck Millipore. The mobile phase was composed of 20 mM ammonium carbonate and 0.1% ammonium hydroxide in water (solvent A), and acetonitrile (solvent B). The flow rate was set at 200 µLl/min with the following gradient: 0 min 80% B, 2 min 80% B, 17 min 20% B, 17.1 min 80% B, and a hold at 80% B for 5 min. Samples were randomized to avoid bias due to machine drift, and the operator was blind to the key. The acquired spectra were analyzed using XCalibur Qual Browser and XCalibur Quan Browser software (Thermo Fisher Scientific) by referencing to an internal library of compounds. Each metabolite is normalised to the total sum of metabolites detected for each sample. ATP was measured enzymatically as described previously [46].

**Soluble protein**

Total protein extraction is described above. Soluble proteins were those that dissolved in reducing denaturing buffer following ethanol precipitation, as follows. 20 µL cell pellets were washed twice in 1 mL ice-cold buffer X, then treated with 200 µl 80°C hot ethanol. The extract was heated for 2 min at 80°C, vigorously mixed on a vortex mixer and incubated for further 2 min at 80°C. The extract clarified by centrifugation, the supernatant removed, and the residual pellet washed 3 times in 1 mL 100% ethanol before being air-dried overnight at room temperature. The dried pellet was resuspended in 200 µL of 8 M urea, 2 M thiourea, 4% CHAPS, 10 mM TCEP and incubated at 37°C with shaking at 400 rpm for 2h to dissolve soluble protein, then clarified by centrifugation at 15,000 rpm for 10 minutes. Soluble and total protein concentration was measured by tryptophan fluorescence (Excitation: 280nm, Emission: 325nm) on a Spark 10M plate reader (Tecan), using BSA to generate a standard curve. Variation in protein content was confirmed qualitatively by SDS-PAGE (Extended Data Fig. 6). Four bands containing soluble protein were excised from denaturing gels, reduced, alkylated and digested with trypsin, using the Janus liquid handling system (PerkinElmer, UK). The digests were subsequently analysed by LC-MS/MS on a Q-Exactive plus orbitrap mass spectrometer, (ThermoScientific, San Jose, USA). LC-MS/MS data were searched against a protein database (UniProt KB) using the Mascot search engine programme [60,61] (Matrix Science, UK). Proteins of the appropriate molecular weight (>70 kD for band 1, 40 to 70kD for band 2, 30 to 40 kD for band 3 and <30 kD for band 4) were included for analysis if they were detected at ≥2% abundance for all three dilution rates. GO annotation and enrichment was carried out using the Saccharomyces Genome Database (SGD).

**Osmolality**

The osmolality of the media was measured using an OSMOMAT 030 (Gonotec), calibrated with 0 and 300 mOsm standards, according to manufacturer's instructions.

**Glycogen**

Cells were harvested by centrifugation, washed with ice-cold PBS and lysed by incubation in 0.25 M sodium carbonate at 98°C for 4 hr. 1M acetic acid and 0.2 M sodium acetate were added to bring the solution to pH 5.2. Glycogen was digested overnight using amyloglucosidase (10115, Sigma-Aldrich) and the glucose released measured using Glucose (GO) Assay Kit (GAGO20, Sigma-Aldrich). Glucose release (mg/ml) was determined from a standard curve and corrected for the OD of starting material.

**Puromycin incorporation**

Samples from the bioreactor were immediately incubated with puromycin (AG Scientific, P-1033) at 8 mg/mL, 30 minutes, 30°C in the dark with agitation, before harvesting by centrifugation. For testing the effect of an osmotic challenge, or change in pH, samples were first diluted into an equal volume of media containing 1% glucose plus glycerol or sorbitol at pH 3.4, or into media of different pH, before addition of puromycin, 8 mg/mL final, 20 minutes.

**Western blot analysis**

Yeast whole-cell extracts were prepared as described [62], except that disruption of the cell wall with glass beads was carried out using a Vortex Genie 2 (Scientific Industries, Bohemia, USA) fitted with a TurboMix head and the protein pellet resuspended in 50 µL Buffer A per OD unit of cells, with protease as [2]. Approximately 10µg protein was separated by gel electrophoresis in MES buffer on 4-12% bis-tris midi NuPAGE gels (Life Technologies) according to the manufacturer's instructions. Proteins were transferred to Immobilon-FL (Millipore) in Towbin buffer using a Biorad Transblot Turbo v1.02 blotter set at 25V, 1.0A, 30 minutes, room temperature, stained for total protein using REVERT (LI-COR) before scanning at 700nm. Blocking was carried out in 1:1 PBS:PBS blocking solution (LI-COR). After incubation with primary and secondary antibodies, the membrane was scanned 800nm and the results quantified using IMAGEJ [63]. The signal intensity of total protein / lane was used for normalisation. All graphs were plotted using Prism. Western blotting using phospho-specific antibodies was carried out using chemiluminescent detection of HRP using TBST containing 0.5% milk and 0.5% BSA.

Primary antibodies: mouse monoclonal anti-puromycin (1:5000, Millipore), mouse monoclonal anti-GFP (1:3000, Roche), polyclonal rabbit anti-Phospho-S6 Ribosomal Protein (Ser235/236), recognises Rps6 in yeast and is a readout for TORC1 activity *in vivo*[64] (1:1000, Cell Signaling), HRP-conjugated antibodies (1:5000, Sigma), were detected using Luminata Forte (EMD Millipore). IR-labelled secondary antibodies were used at a 1:10,000. Blots were scanned on a gel doc (Biorad) or Odyssey FC (LI-COR).

**Protein aggregation**

Aggregated protein was prepared as described [65]. Proteins were visualized by silver staining using the Silver Quest Kit (Invitrogen) and images captured using a Perfection V600 Photo scanner (Epson). ImageJ was used to obtain the background corrected signal intensity per lane [63].

**Flow cytometry**

Propidium iodide labelling was carried out as previously [2] and samples processed at the CCTI Flow Cytometry Core on a FACS Canto II (Becton Dickinson). Analysis was performed using the FCS Express density plot function, with the gates shown in Extended Data Fig. 1.

**Intracellular pH**

***Ratiometric bioluminescence monitoring during the YRO***

YROs were established in strains with a stably transformed bioluminescent pH reporter plasmid (either pRS306-PTEF1luc2 or pRS304Nat- PTEF1luc2), 850 ml culture in a 3L BioFlo 115 benchtop fermentor (New Brunswick) with pH 3.4, 54 L/h air, 550 rpm, and an initial dilution rate of 0.08/h [66]. Luminescence was continuously recorded using two Hamamatsu PMTs (HC135-01) [67]; however, the PMTs were fitted with 550 ±5-nm and 610 ± 5-nm band pass filters (65704 and 65709; Edmund Optics).

***pH – Luminescence ratio curve***

After monitoring luminescence from the YRO (as above), continuous culture was terminated. The pH of the culture was adjusted to 5.8 by addition of 2M NaOH. Beetle luciferin (Promega) was added (5 µM final). And carbonyl cyanide m-chlorophenyl hydrazone (CCCP, Promega) was added (1.18 µM final). The culture incubated for 4h. The pH was increased stepwise by 0.2 by addition of 2M NaOH every 15 min covering a pH range 5.8 – 8.2. Bioluminescence at 550 and 610nm was continuously monitored as above.

**Confocal microscopy**

Samples from the bioreactor were incubated in the dark with 7-amino-4-chloromethylcoumarin (CMAC, LifeTech C2110) 20µM (final), 30°C, 10 minutes with agitation. Cells were washed 1x in media without glucose before imaging.

All microscopy was carried out using an Axioskop2 (Zeiss) equipped with a 100x/1.4 Plan-Apochromat objective, Orca ER cooled charge-coupled device (CCD) camera (Hamamatsu) and pE-4000 CoolLED light source at 405 nm, 200 ms for CMAC and 470 nm, 250 ms for GFP (21 image *z* stack at 0.3 µm). Images were deconvolved using iterative restoration and analysed using Volocity 5.3 and 6.5 (Perkin-Elmer) using default settings.

**Survival after heat stress**

Samples from the bioreactor were divided into two aliquots. 150µL was incubated at 55°C for 2 minutes before serial dilution in media without glucose. The other was diluted in parallel and plated directly (untreated control). The number of colonies/plate was counted after 2 days at 30$^o$C (median cells per plate 279) and number of colonies on heat treated vs. non-treated samples expressed as a percentage.

**Osmotic challenge**

Potassium chloride (3M in media without glucose) was added to 200mM final concentration. Sodium chloride (s), to 1M final concentration, was added directly to cells in the bioreactor.

**Statistics**

Unless otherwise stated, statistical analyses were performed using Prism 8 (Graphpad). Grubbs' method for outlier detection was employed with 〈=0.0001 for every species detected by ICP-MS and LC-MS, and found a single consistent outlier (replicate 1, time point 5, dilution rate 0.06$h^{-1}$) with values that were erroneously high. This was excluded from subsequent analysis.

**Data and code availability**

All data and code are available on request.

**Yeast strains**

All strains are isogenic *Mata* haploids in the CEN.PK113-7D background unless stated.
HCY1514 *prototrophic wild type*
HCY1602 *HSP30::KanMX*
HCY1648 *GPH1::KanMX*
HCY1674 *GSY2::KanMX*
HCY1325 *Mata/Matα*
HCY1689 to HCY1691 *Mata/Matα PGK1/PGK1*-GFP-KanMX
HCY1800 *Mata* yBR-PTEF1luc2
HCY1803 *Mata* BR-PTEF1luc2 *HSP30::KanMX*
HCY1820 *Mata ura3Δ0* [pRP1657 Pab1-GFP Edc3-mCherry] [68]

Gene inactivation was carried out by amplification of the KanMX antibiotic resistance cassette from either the pFA6 plasmid [69], or another deletion strain and integrated into the genome of a CEN.PK strain. The location of the cassette was confirmed by PCR and/or sequencing.

***Reporter for autophagy***

Pgk1/Pgk1-GFP-KanMX strains were made by amplification of the GFP-KanMX cassette from pFA6 plasmid[69] using primers:
5'-TAAGGAATTGCCAGGTGTTGCTTTCTTATCCGAAAAGAAACGGATCCCCGGGTTAATTAA-3'
5'-CTTAAAATACGCTGAACCCGAACATAGAAATATCGAATGGGAATTCGAGCTCGTTTAAAC-3' generating an amplicon containing the end of *PGK1*, a 4 amino acid spacer, GFP, KanMX and the 3' UTR of *PGK1*. This was integrated into the genome of diploid CEN.PK at a single locus. Cleavage of Pgk1-GFP fusion protein by non-selective bulk autophagy can be detected by western blotting [70].

***Construction of strain containing a bioluminescent reporter of intracellular pH***

pRS306ΔXbaI was created by removing the XbaI site in pRS306 [71] by digesting the plasmid with XbaI, treating the digested product with DNA polymerase I large fragment and recircularizing. The BamHI-XhoI fragment from pRS306-PTEF1CBR [67] containing the *TEF1* promoter, CBR luciferase, and *ADH1* terminator was transferred to the pRS306ΔXbaI plasmid to produce pRS306-PTEF1CBRΔXba. The *luc2* CDS was PCR amplified from pGL4.20 using primers:
5'-actactAGATCTATGGAAGATGCCAAAAACATTA-3',
5'-actacgTCTAGATTATTTTTCGAACTGCGGGT-3' adding a BglII and XbaI site upstream and downstream of the start and stop codons respectively. Finally, the *luc2* PCR product was added to pRS306-PTEF1CBRΔXba with BglII and XbaI (replacing the CBR luciferase CDS) and generating pRS306-PTEF1luc2, an integrating plasmid for constitutive *luc2* expression. pRS306-PTEF1luc2 was linearized with EcoRV and transformed into yBR-ura3ΔCEN.PK113-7D [72], creating strain yBR-PTEF1luc2.

A nourseothricin selectable bioluminescent pH reporter was made by putting the BamHI-SalI fragment from pRS306-PTEF1luc2 (containing the *TEF1* promoter, luc2 luciferase, and *ADH1* terminator) into pRS304-Nat [67] creating pRS304Nat- PTEF1luc2. This plasmid was linearized with EcoRV and integrated into the native *TRP1* gene of strain HYC1602.

### Extended Data Figure 1

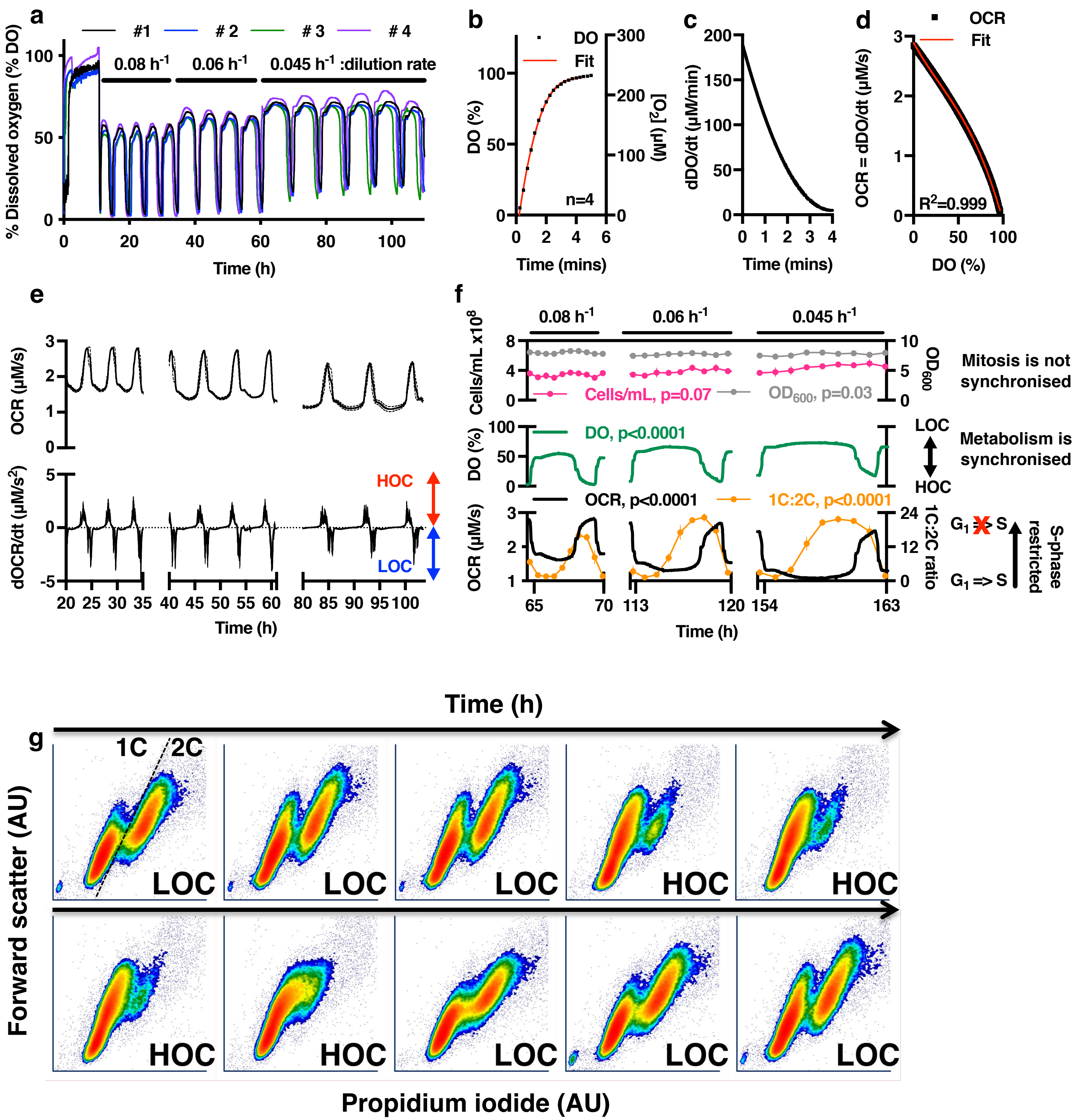

### Extended Data Figure 2

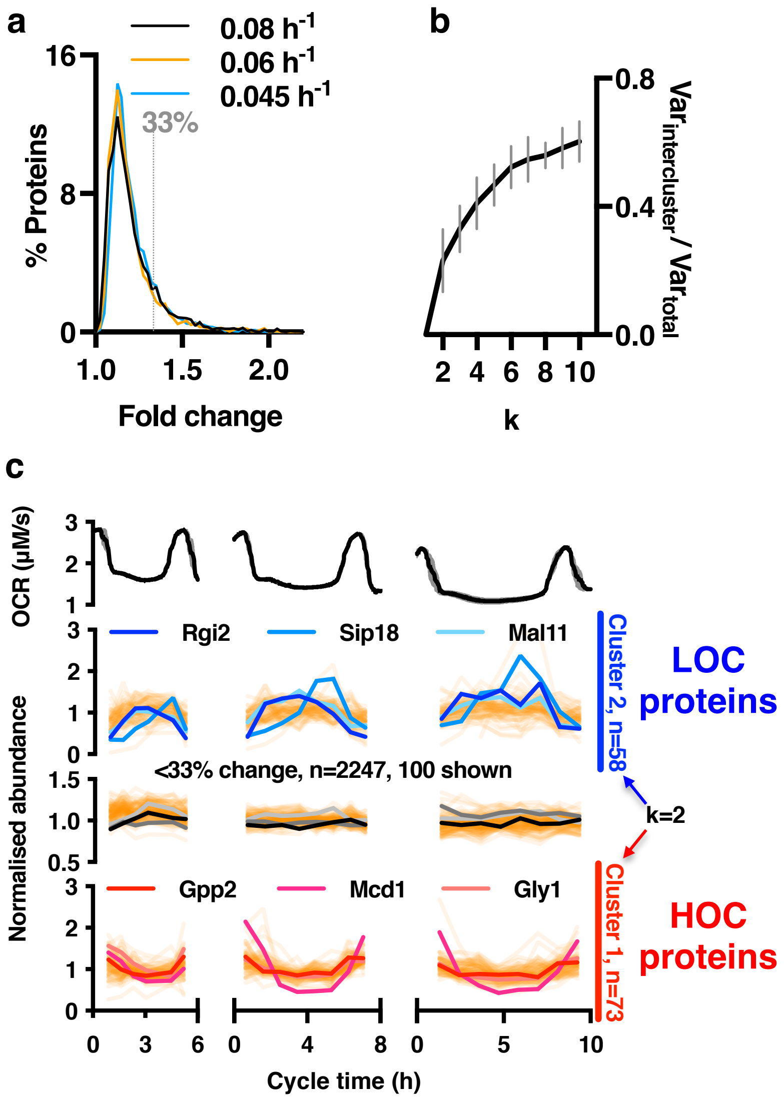


d

| GO term | Description | P-value | Enrichment | Gene name |
|---|---|---|---|---|
| HOC enriched process (>33% amplitude) | | | | |
| 0072348 | sulfur compound transport | 0.00015 | 23.76 | SUL1 VHT1 THI7 |
| 0055085 | transmembrane transport | 0.00044 | 3.03 | SUL1, VHT1, THI7, ENA1 HIP1, AGP1 FCY2, MGR2, TIM8, YMR166C, YOR020W-A, PEX25 |
| 0006114 | glycerol biosynthetic process | 0.00063 | 39.61 | RHR2, HOR2 |
| LOC enriched process (>33% amplitude) | | | | |
| 0015718 | monocarboxylic acid transport | 0.00002 | 21.66 | FMP43, JEN1 ATO2, ATO3 |
| 0010035 | response to inorganic substance | 0.00010 | 10.15 | NCE103 CUP1-2, CRS5 SIP18, STF2 |

### Extended Data Figure 3

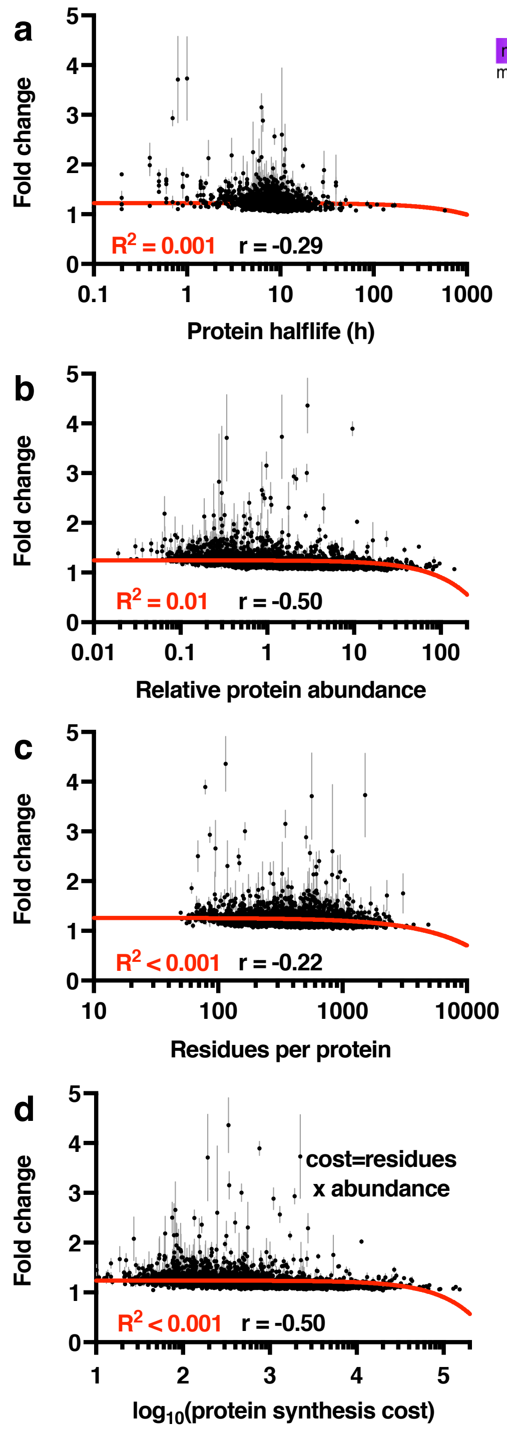


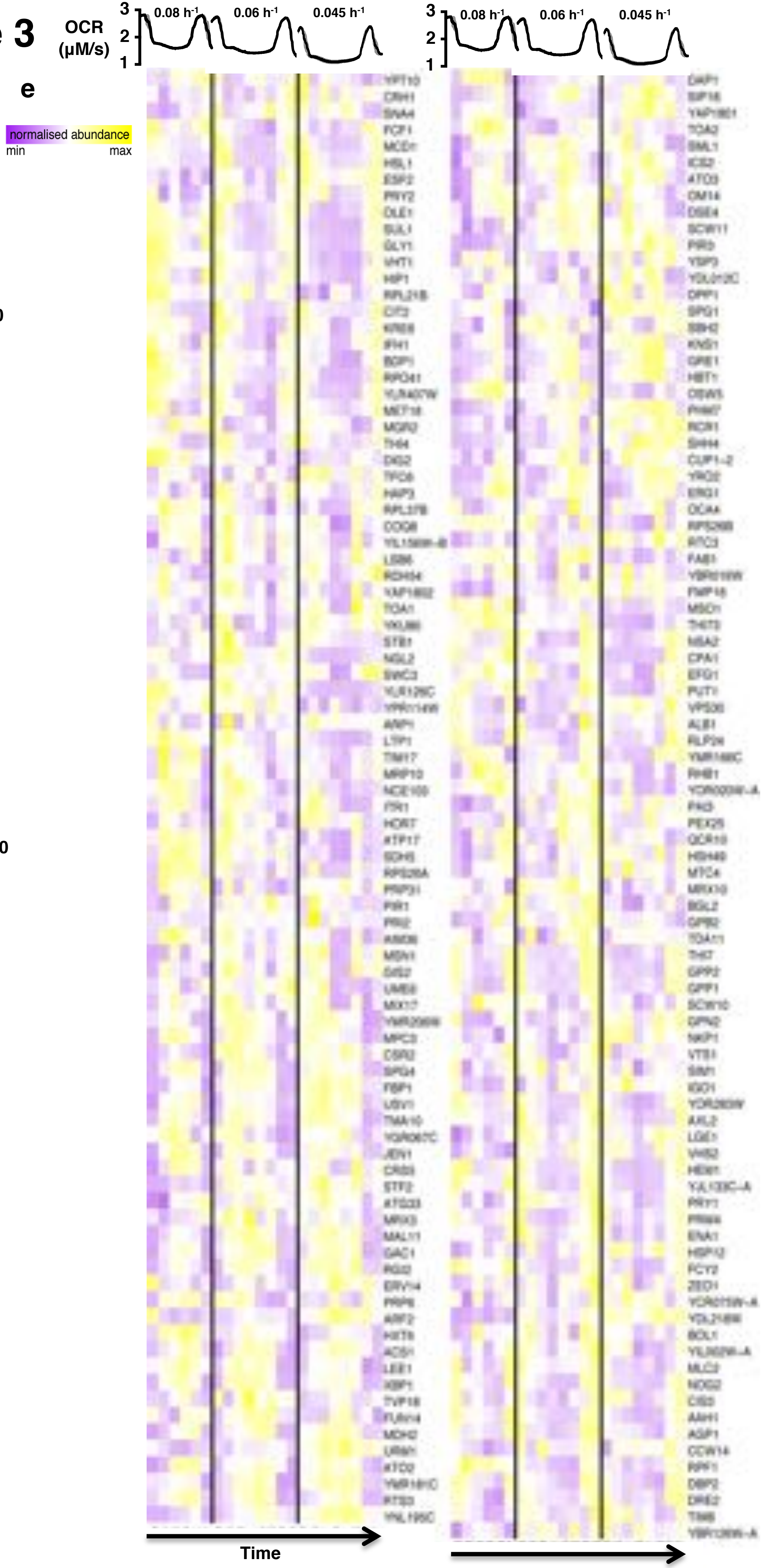

### Extended Figure 4

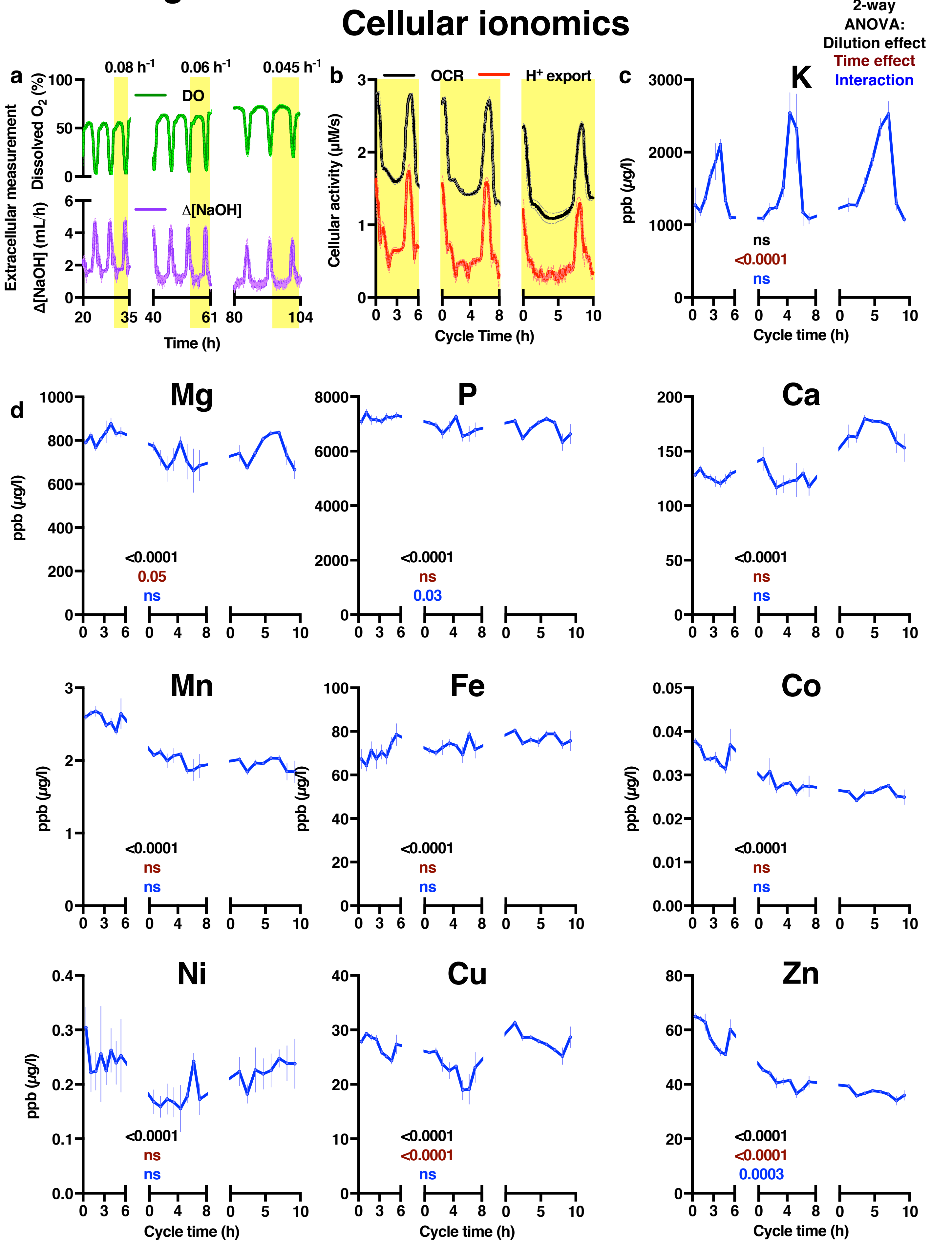

### Extended Data Figure 5(i)

#### Cellular metabolomics

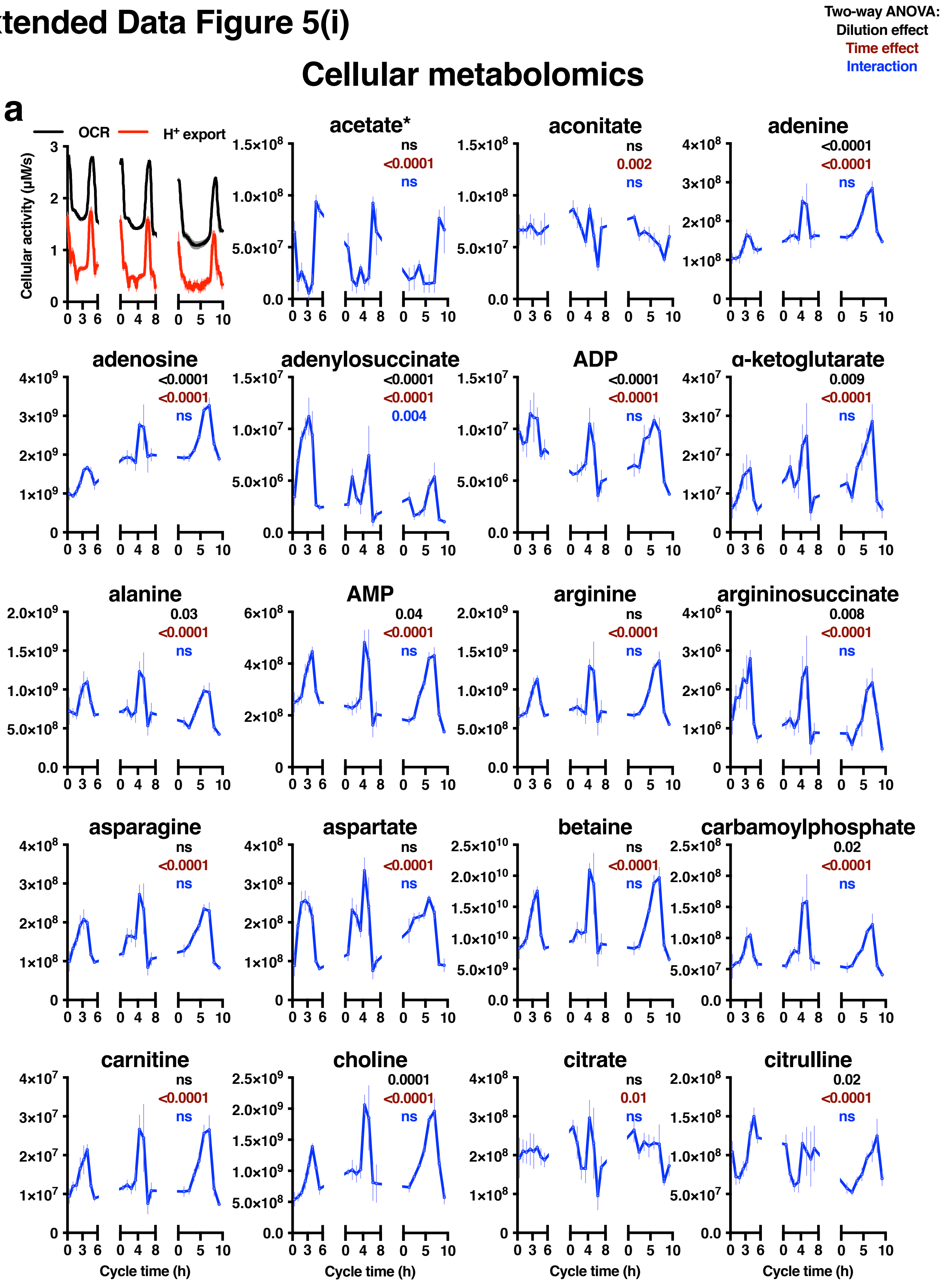

### Extended Data Figure 5(ii)

Two-way ANOVA:
Dilution effect
Time effect
Interaction

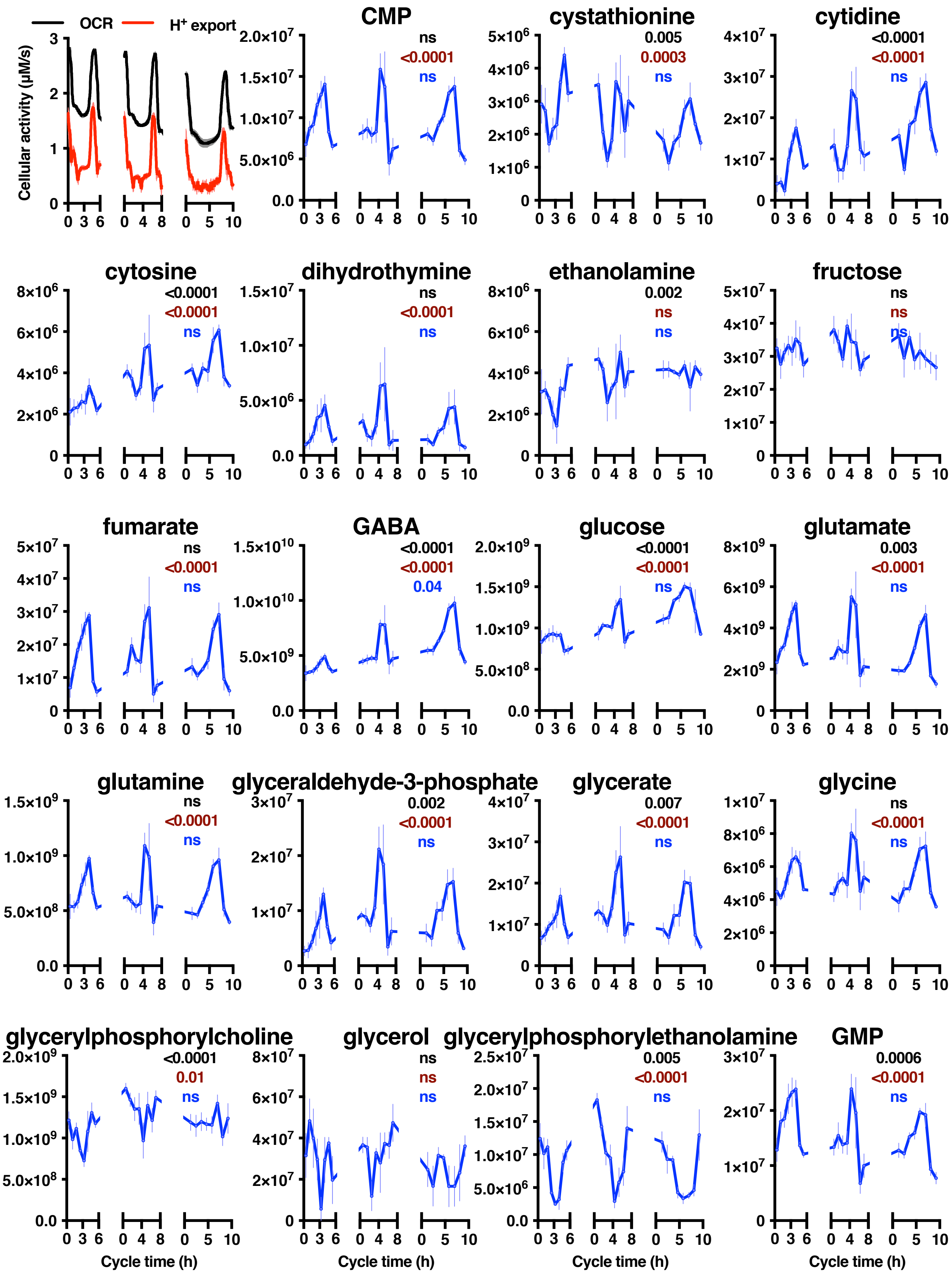

### Extended Data Figure 5(iii)

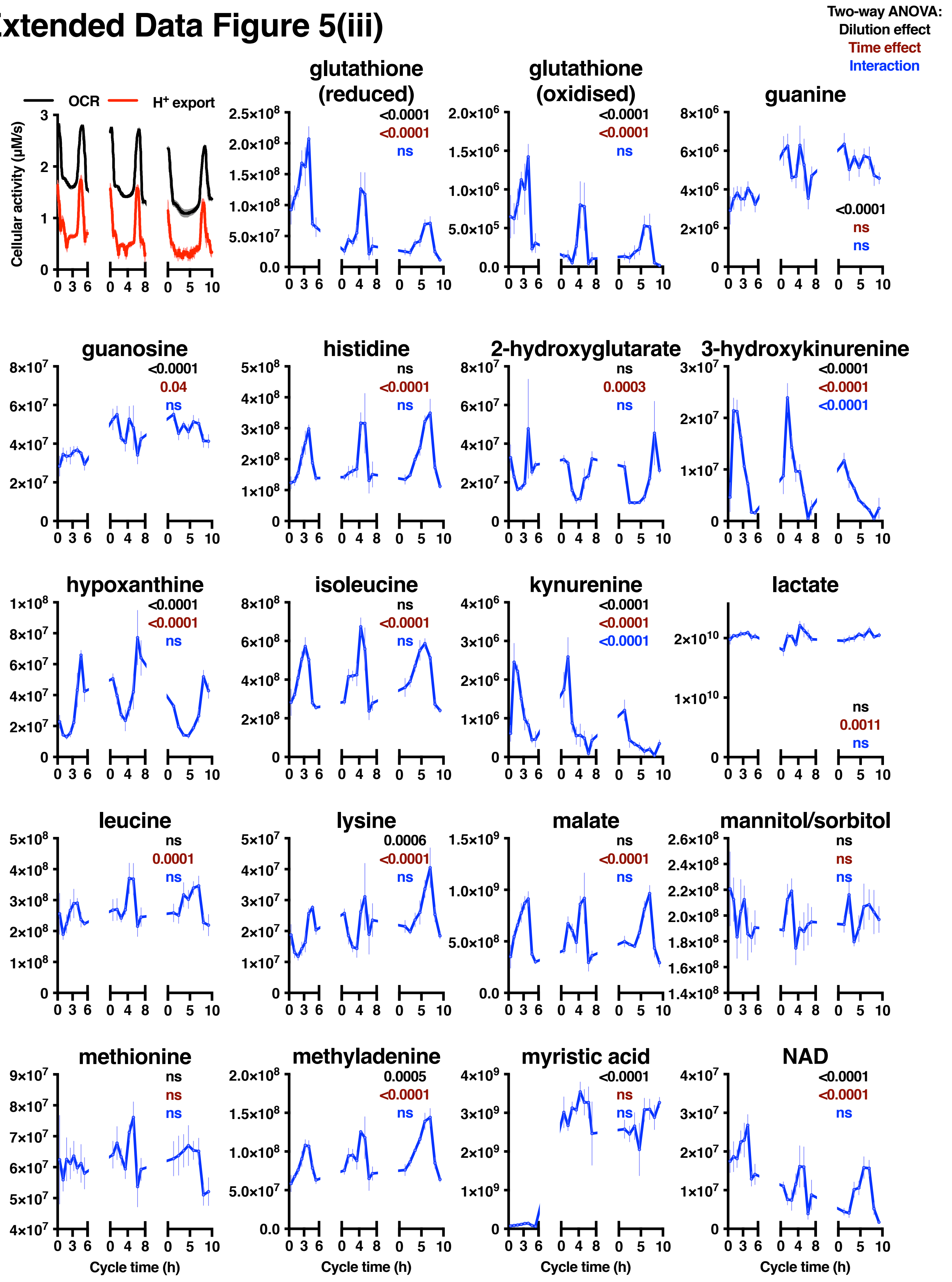

### Extended Data Figure 5(iv)

Two-way ANOVA:
Dilution effect
Time effect
Interaction

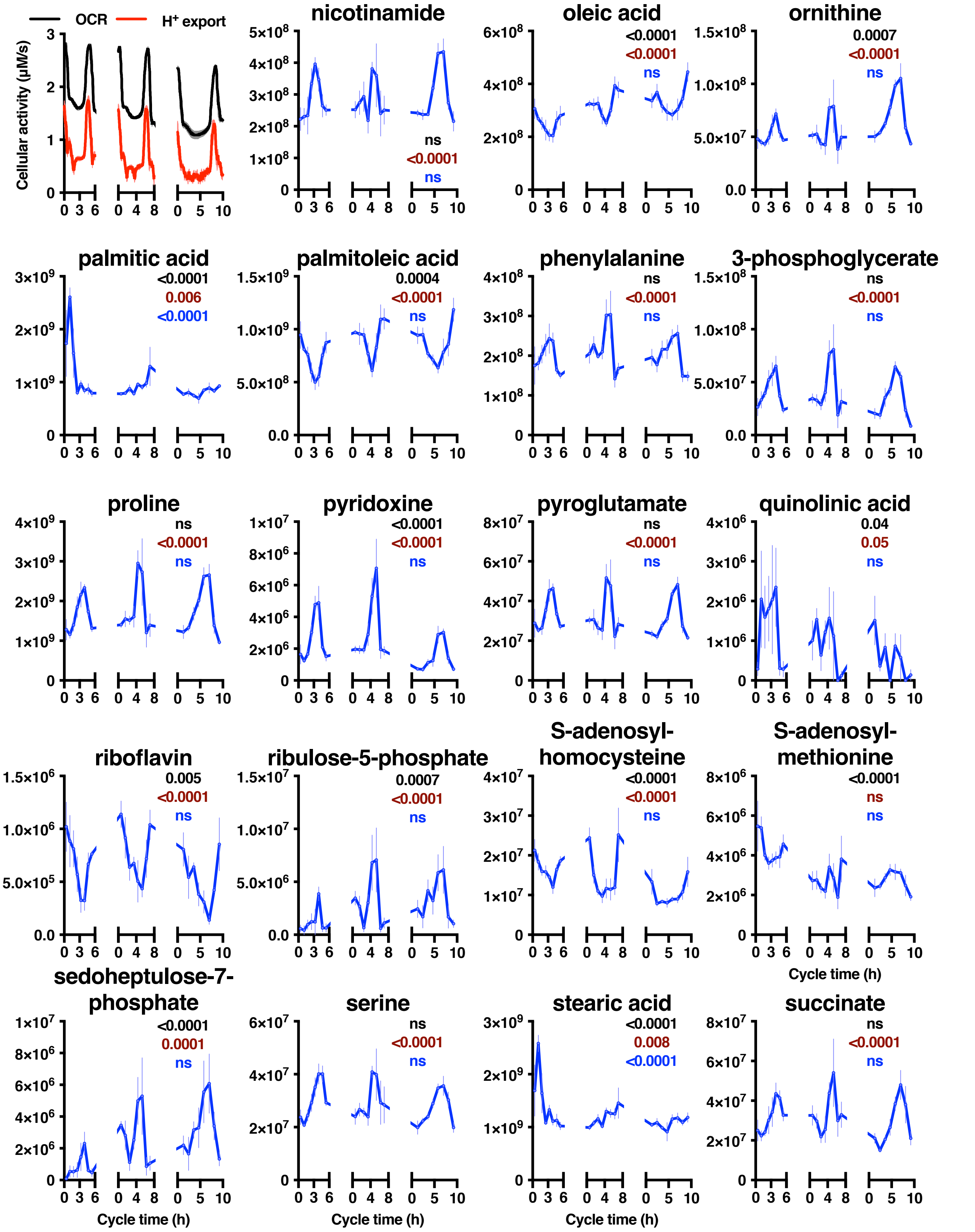

### Extended Data Figure 5(v)

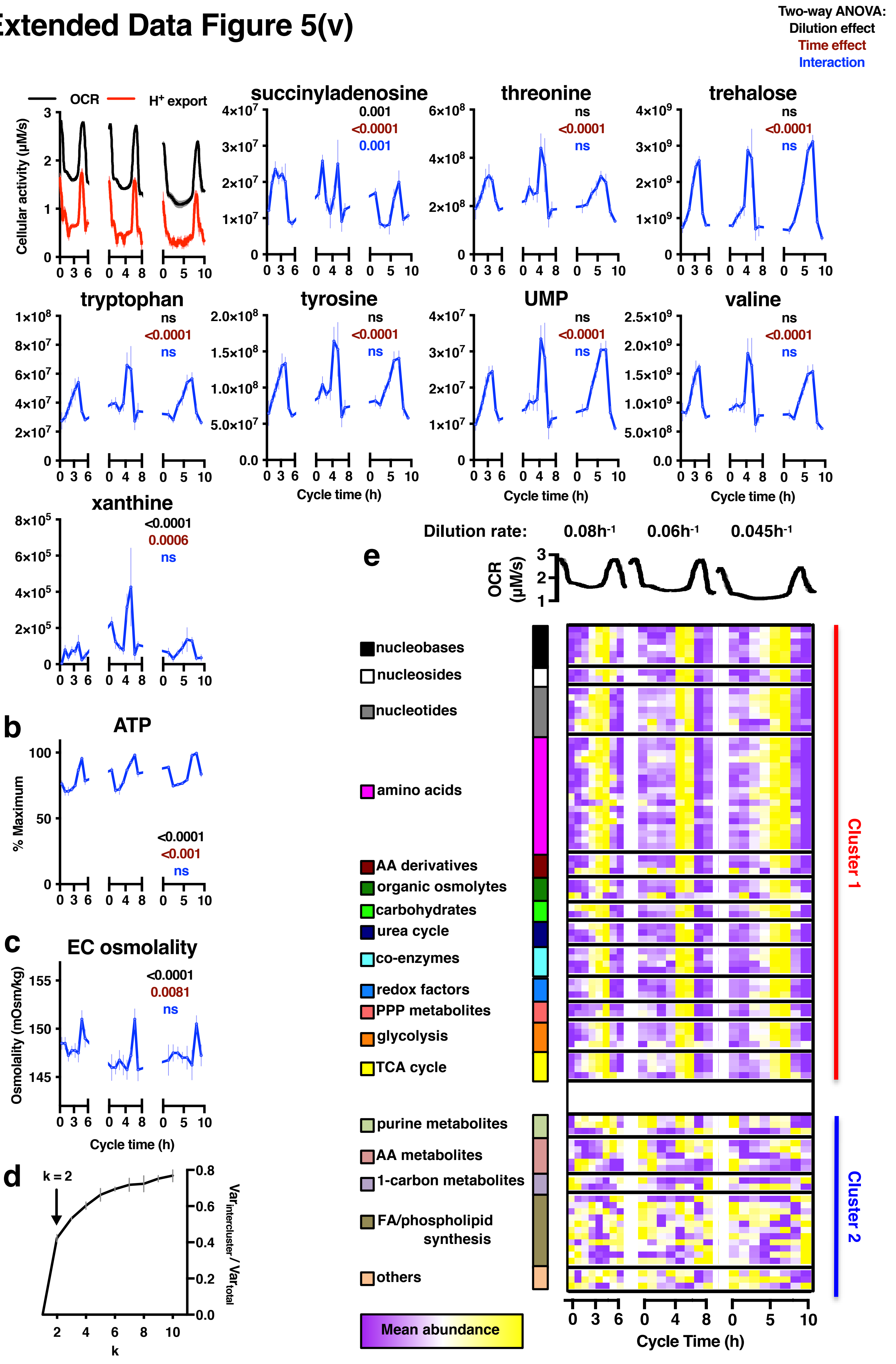

### Extended Data Figure 6

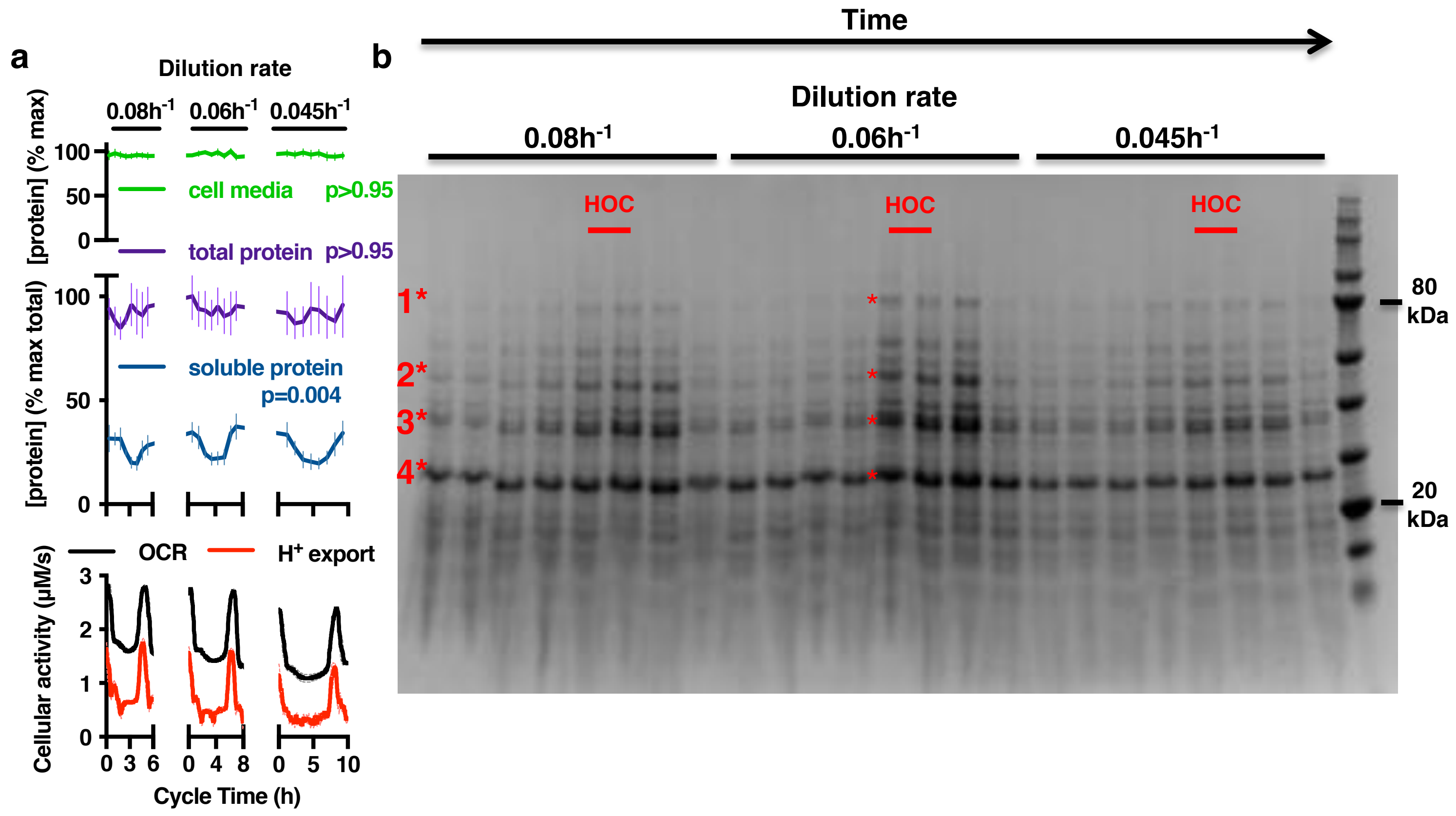


c

| Band | Enriched for GO term | Proteins with GO annotation |
|---|---|---|
| 1 | Protein-binding involved in protein folding, p = 2.06 x $10^{-13}$ | Ssa2, Ssa3, Ssc1, Kar2, Ssa1, Ssa4 |
| 2 | Catalytic activity, p = 4.56 x $10^{-11}$ | Atp2, Erg6, Arc1, Met17, Pdx1, Err3, Eno2, Arg1, Idp3, Hsp60, Eno1, Cit1, Pot1, Mri1, Pmi40, Sam1, Pda1, Gdh1, Kgd2, Pgk1, Tef2, Ald4, Tuf1, Idp1, Ddi1, Lys9, Cor1, Sah1, Pdc1, Hat2, Sam2, Rpt6, Oye3, Cdc19, Cpr6 |
| 3 | Oxidoreductase activity, p = 3.63 x $10^{-14}$ | Mtd1, Mdh1, Adh2, Zta1, Tdh2, Gre3, Hom6, Ypr1, Mrp1, Tdh3, Gcy1, Tdh1, Ara1, Aim45, Ydl124w, Lia1, Ypr127w, Adh1 |
| 4 | Proteasome/endopeptidase activity, p = 2.04 x $10^{-6}$ | Pre9, Pre8, Scl1, Pre7, Pup2 |
| 4* | 40S ribosomal subunit, ns (p = 0.06) | Rps3, Rps5, Rps0b, Rps8b |

### Extended Data Figure 7

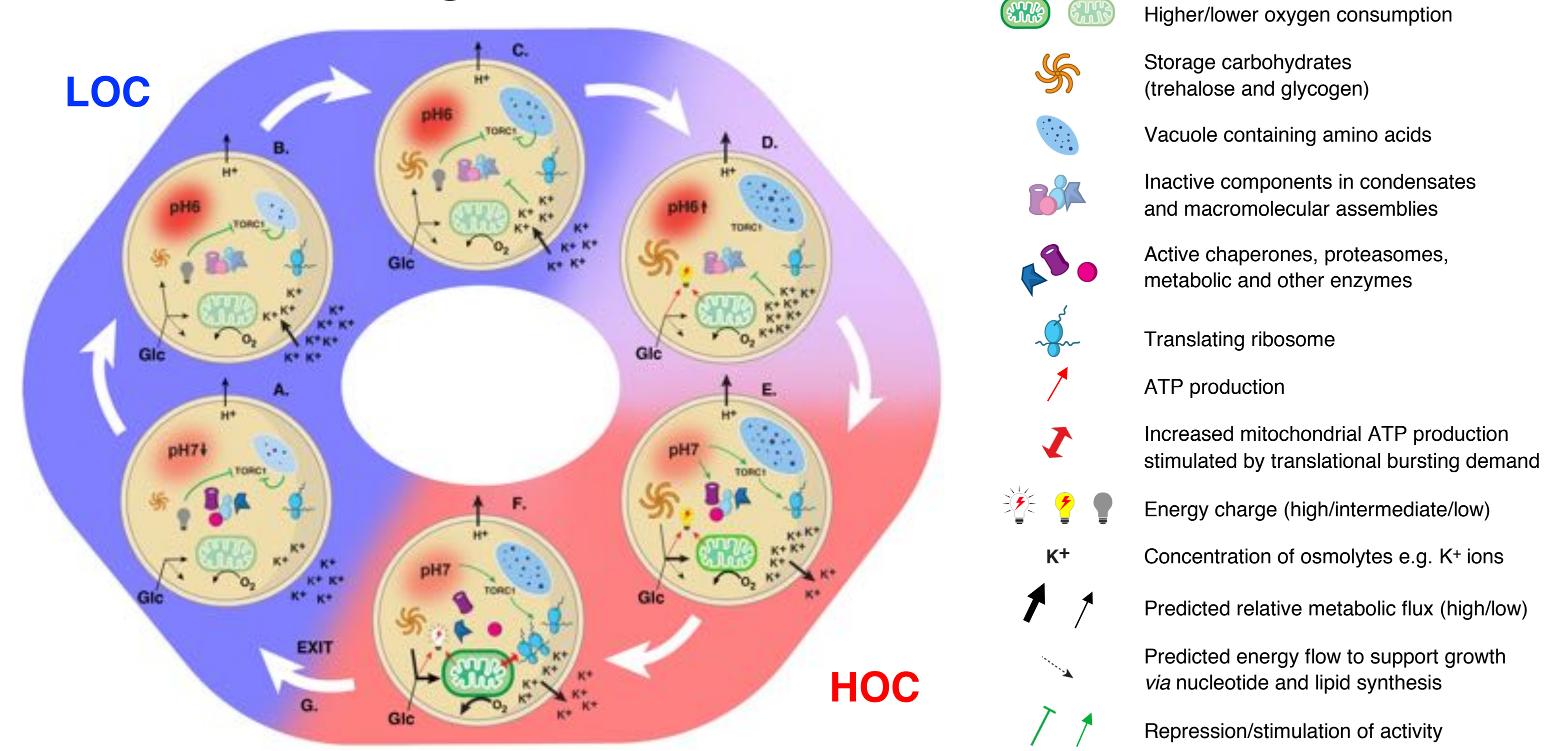


**Extended Data Figure 7, A detailed, testable and experimentally-derived model for the YRO.**
During exponential growth, glucose is not limiting. Protein synthesis and growth (biomass accumulation, DNA synthesis, fatty acid synthesis) do not compete for bioenergetic resources.

**A. Early LOC**

- Storage carbohydrates and amino acids are depleted, glucose supply is low (Fig.2a, [11,16,23,73-78]).
- Low glucose inhibits Pma1, the ATP-dependent plasma membrane $H^+$-pump in yeast, resulting in a low rate of $H^+$-export [19,20,43,79-83].
- Cytosolic pH drops (Fig.2c, Ext.Fig.4a), coordinately regulating the activity of many metabolic and signalling pathways [84] *e.g.* inhibition of glycolysis [85].
- Low cytosolic pH and glucose availability leads to reduced glycolytic flux & $O_2$ consumption, with decreased rates of ATP production[11,86] eliciting a fall in energy charge (Fig.2a,c, Ext. Fig.5).
- TORC1 is inactivated (Fig.3b) due to: (1) low energy charge *via* SNF1/AMPK (Fig.2a, Ext. Fig.5 [41]); (2) amino acid depletion (Fig.2a, Ext. Fig.5) *via* Gcn2 [40]; (3) inhibition of the activating GTPase Gtr1/2 due to low pH [22,87].
- TORC1 inactivation results in decreased protein synthesis (Fig.3b, [40]) and stimulates autophagy (Fig.3c, [88]).

**B&C. Mid-Late LOC**

- Pma1 activity is further limited by reduced ATP availability [19,43,84].
- Cytosol stabilizes at ~pH6.3, initiating a cellular response to starvation stress which facilitates macromolecular assembly of cytosolic enzymes and increased formation of biomolecular condensates (BMCs) such as stress granules and p-bodies (Fig.2a, Fig.4a, [24,25,27,83,89-96]); resembling a quiescent state where stress-resistance is enhanced and the cytoplasm is viscous and more 'glass-like' (Fig.5b; [31,74,97,98,101,102]).*
- TORC1 inactive (Fig.3b), reduced protein synthesis (Fig.3b), increased autophagy (Fig.3c), low oxygen consumption (Fig.2a) and energy charge (Fig.2a, Ext.Fig.5).
- Sequestration of cytosolic metabolic enzymes such as Cdc19[27] (Ext. Fig.6b,c), low pH and reduced glycolytic flux [85] direct glucose to production of storage carbohydrate (glycogen and trehalose, Fig.2a, [11,23,36,74,77,78,94]), and generate biosynthetic intermediates for cell growth *via* the pentose phosphate pathway, fatty acid and DNA synthesis (Ext.Fig1, Ext.Fig5, [99]).
- The ~1000-fold $H^+$ gradient (pH 3.4 extracellularly) across the plasma membrane is used in secondary active transport to accumulate nutrients and osmolytes (Fig.2a,b, [19,100]).
- Import of $K^+$ and other osmolytes (Fig.2a, Ext.Fig.4c) counter-balances the reduced contribution that sequestered cytosolic macromolecules make to cellular osmotic potential [32,100].
- Autophagy and amino acid symporters act to replenish vacuolar amino acid stores [103,104].

**D. Entry to HOC**

- Replete carbohydrate stores (Fig.2a, 4b) stimulate glycolysis, respiration and ATP production [105,106].
- Increased ATP availability and higher glycolytic flux increase Pma1 activity [19,43,84] and so cytosolic pH begins to increase (Fig.2a,c).
- Energy charge increases due to increased ATP production relative to consumption and associated fall in AMP/ADP (Fig.2a, Ext. Fig.5).
- Increased energy charge relieves Snf1/AMPK-mediated inhibition of TORC1 [41] and stimulates glycogenolysis [105].
- Replete amino acid stores relieve Gcn2-mediated inhibition of TORC1 [40].
- Condensate disassembly is initially attenuated by high cytosolic osmolyte concentration ($K^+$, choline, betaine) and low pH (Fig.2a,c; Ext. Fig.4,5).

**E. Early HOC**

- Elevated cytosolic pH and increasing energy charge trigger the release of sequestered ribosomes, proteasomes, chaperones and metabolic enzymes such as Cdc19 from macromolecular assemblies and condensates [24,25,27,89-95,107] – a feed-forward switch that further stimulates glycolysis (Ext Fig.5) and glycogen/trehalose breakdown (Fig.4b, Ext.Fig.5, [108]). This further increases ATP production and the rate of $H^+$-export (Fig.2a,c, [43,81,109]). Cytosolic pH reaches pH7 (Fig.2c).
- Increased cytosolic pH activates TORC1 [22,87] which stimulates increased translational initiation and represses autophagy [38,40,88], preventing futile cycles.
- To maintain osmotic homeostasis, osmolytes are exported down their concentration gradients. This buffers the increase in osmotic potential due to increased cytosolic macromolecules, and the cytosol becomes more fluid [31,32,97,100-102].

**F. Late HOC**

- Glycogen and trehalose breakdown sustain high glycolytic flux and respiration (Fig.2a, 4b, Ext. Fig.5).
- Increased mitochondrial ATP production rate is stimulated by, and sustains, the high rate of ATP turnover required for efficient protein synthesis [110,111]. The proportion of ribosomes available for translation much higher than during LOC (Ext. Fig.6, [31]).
- Continued export of osmolytes counter-balances the osmotic potential due to newly synthesized macromolecules [31,32,100].
- Increased protein synthesis consumes stored amino acids (Fig.2a, ExtFig.5).
- DNA replication and fatty acid synthesis do not occur (Ext. Fig.1f,g) as all cellular glucose is used to sustain translational bursting, which now consumes up to 75% of cellular energy [28,110,112].

**G. HOC Exit**

- Decreased protein synthesis leads to decreased demand for ATP turnover and so oxygen consumption falls (Fig.2a, 3b, Ext.Fig.5b). This could be due to insufficient: stored osmolytes to buffer further protein synthesis (equivalent to osmotic stress, Fig.2a, Ext.Fig.5 [113]); stored amino acids to sustain further protein synthesis [40]; stored carbohydrates or oxygen to meet the requirements of protein synthesis and $H^+$-export (*via* TORC1, [22,41,87]).
- This predicts that premature HOC exit will occur on acute osmotic stress (Ext. Fig.8b), inhibition of protein synthesis or TORC1 activity (Fig.3e), and that perturbation of osmotic buffering capacity, the transmembrane $H^+$-gradient, or Pma1 activity will alter the period of oscillation (Fig. 4c-h).

*The mechanisms that facilitate widespread macromolecular assembly formation and condensation in low glucose are incompletely understood, but have been widely observed, and are likely to involve increased association of proteins with RNA, and other proteins, through electrostatic charge-charge interactions made favourable by histidine protonation, lysine/arginine modification and changes in protein phosphorylation, as well as changes in ionic strength and the chemical potential of water [25,114-121].

### Extended Data Figure 8

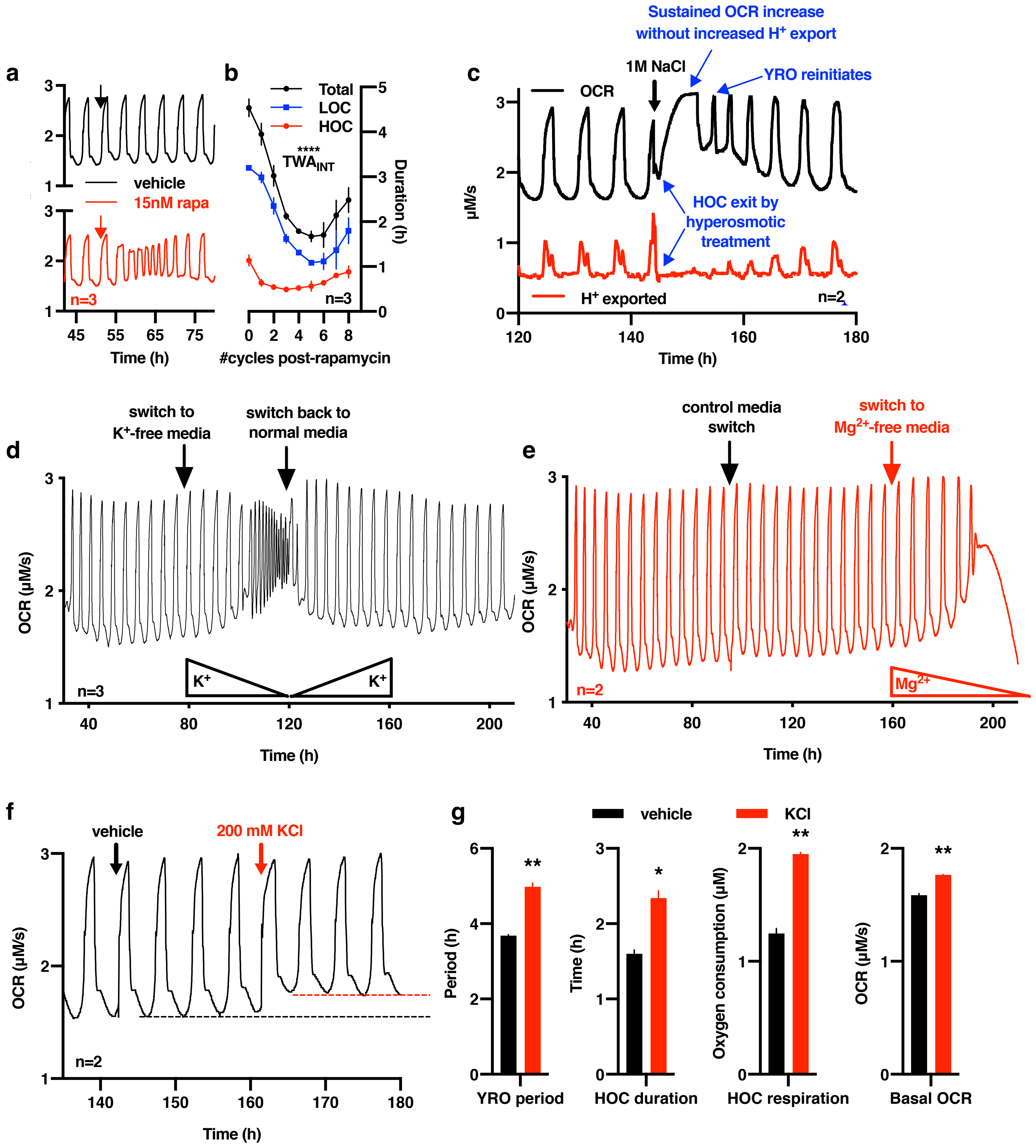

### Extended Figure 9

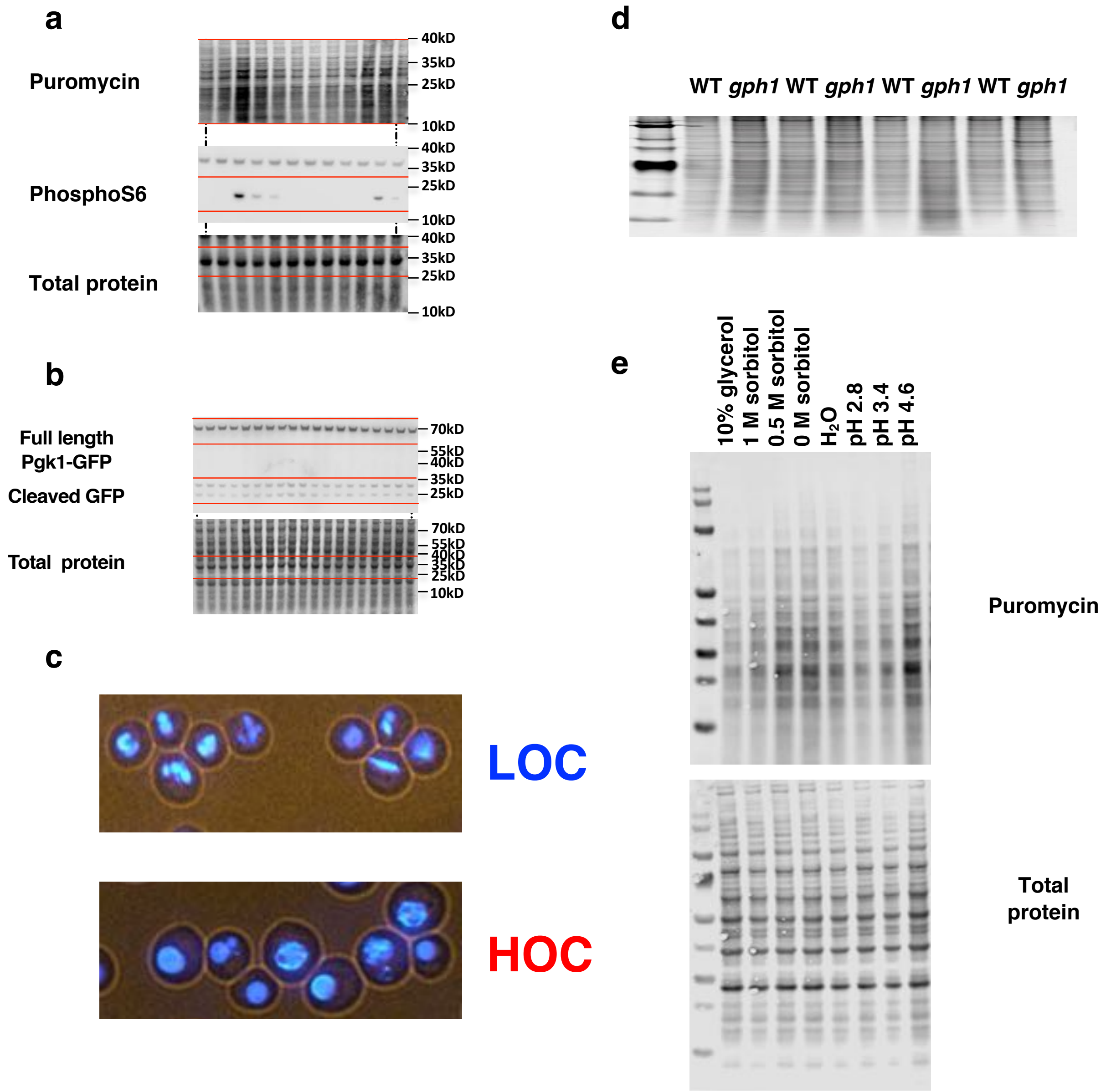
